# OCRL regulates lysosomal function and endolysosomal homeostasis in *Drosophila* nephrocytes

**DOI:** 10.64898/2025.12.03.692239

**Authors:** Vaishali Kataria, Navyashree A Ramesh, Indra Sara Lama, Aishwarya Venugopal, Anil Vasudevan, Padinjat Raghu

## Abstract

The *OCRL* gene encodes a lipid phosphatase that dephosphorylates phosphatidylinositol 4,5 bisphosphate [(PI(4,5)P_2_]. Mutations in *OCRL* lead to a rare human genetic disorder, Lowe syndrome (LS) that affects the eye, kidney and brain. OCRL is widely expressed in cells and is localized to multiple organelles, including the plasma membrane, endosomes, Golgi and lysosomes. Although multiple defects in the endo-lysosomal system have been reported in *OCRL* depleted cells, the primary site of action of OCRL is unclear. Here we present a *Drosophila* nephrocyte model of LS; depletion of *Drosophila* OCRL (*dOCRL*) manifests with defects in endocytic uptake, altered endosomal compartments as well as expanded but dysfunctional lysosomes. Reconstitution of *dOCRL* depleted nephrocytes with a lysosome targeted version of the enzyme rescues not only the lysosomal defects but surprisingly also defects in endosomal structure and function. These findings suggest that the primary defect in LS cells is likely to be altered lysosome structure and function. Therefore, regulation of PI(4,5)P_2_ homeostasis at the lysosome membrane by OCRL is critical to homeostasis of the endosomal system.

## Introduction

Lowe syndrome (LS) is a rare X-linked recessive genetic disorder characterized by clinical features of renal tubular dysfunction, mental retardation and cataract (De Matteis et al., 2017; Preston et al., 2020). The disease results from the mutations in the *<u>o</u>*culo*<u>c</u>*erebro*r*enal syndrome of *<u>L</u>*owe (*OCRL)* gene that encodes a type II family inositol polyphosphate 5-phosphatase enzyme involved in the hydrolysis of phosphate at 5^th^ position of phosphatidylinositol-4,5-bisphosphate [PI(4,5)P_2_] to generate phosphatidylinositol 4 phosphate (PI4P) (Attree et al., 1992; Zhang et al., 1995). OCRL is a multi-domain enzyme with a catalytic 5-phosphatase domain, N-terminal pleckstrin homology (PH) domain, an ASPM-SPD-2-Hydin (ASH) domain and a Rho GTPase-Activating Protein (RhoGAP) domain (De Matteis et al., 2017). The *OCRL* gene is widely expressed across human cells and tissues and throughout development; the protein is localized to multiple sub-cellular membranes in human cells including the plasma membrane, endolysosomal compartments, trans-Golgi, cilium and lysosomes [reviewed in (Mehta et al., 2014)].

Proximal tubular dysfunction in the kidney is a key clinical manifestation of LS, characterized by renal tubular acidosis, low molecular weight proteinuria, hypercalciuria, albuminuria, aminoaciduria, carnitine wasting, phosphaturia with perturbed glomerular filtrate rate that deteriorates with age (Bockenhauer et al., 2008; Recker et al., 2015). Several studies have attempted to decipher the sub-cellular origin of altered proximal tubular function in LS. Studies in COS-7 (kidney derived) cells found OCRL localized to early endosomes and the Golgi apparatus (Ungewickell et al., 2004) and early endosomal function was impaired by OCRL depletion (Vicinanza et al., 2011). This altered endosomal maturation was proposed to result from abnormal actin polymerization on early endosomes (Nández et al., 2014; Vicinanza et al., 2011). Studies in a zebrafish model of LS have reported alterations in the endosomal system (Oltrabella et al., 2015) and this has also been reported in a humanized mouse model of LS (Festa et al., 2019). A *Drosophila* model of LS has reported alterations in the endosomal system of haemocytes and proposed to affect other tissues in a non-cell autonomous manner (Del Signore et al., 2017). On the other hand, studies with human kidney biopsy samples have reported OCRL localized on lysosomes (Zhang et al., 1998) and a role for OCRL in regulating autophagosome-lysosomal function has been proposed (De Leo et al., 2016). Indeed an early study had reported increased levels of several lysosomal enzymes in the circulation of LS patients (Ungewickell and Majerus, 1999). Thus, despite multiple studies reporting sub-cellular defects in the endo-lysosomal system on OCRL depletion, it is unclear whether OCRL regulates function at each of these compartments individually or whether there is a primary site of action of OCRL within the endosomal system and the defects in other compartments represents a cellular response to maintain homeostasis in the endosomal system.

The filtration of blood in the glomerulus of nephrons followed by selective reabsorption of essential molecules is a key aspect of human kidney function. In *Drosophila* too, nephrocytes perform an analogous function to purify circulating haemolymph. *Drosophila* larvae have two types of nephrocytes, garland and pericardial, tethered to the oesophagus and heart respectively. Nephrocytes take up haemolymph by endocytosis, sort the endocytosed cargo for degradation in lysosomes or recycle it back to the cell surface. At a molecular level, the function of the multi-ligand receptor megalin in proximal tubule cells (PTC) mediates protein reabsorption and its trafficking is affected by OCRL depletion (Oltrabella et al., 2015; Vicinanza et al., 2011). In *Drosophila*, a functional ortholog of megalin, the cubulin/amnionless co-receptor (Zhang et al., 2013) has been described in the fly nephrocyte. Thus both endocytic function as well as lysosomal degradation of endocytic cargo is central to *Drosophila* nephrocyte function. The endocytic function of fly nephrocytes is similar to that seen in proximal renal tubule cells and hence offers an opportunity to model various aspects of proximal tubular functions (Helmstädter et al., 2017; Koehler and Huber, 2023). Owing to their large size, *Drosophila* nephrocytes allow both live cell imaging of organelle structure, function, and analysis of physiological readouts from intact animals.

In contrast to human genome, the *Drosophila* genome contains a single homolog of *OCRL* (Balakrishnan et al., 2015; Ben El Kadhi et al., 2011) and therefore offers an opportunity to analyse function without the complexity of contributions from multiple genes (e.g *OCRL* and *INPP5B* in the human genome) that encode the same inositol polyphosphate 5-phosphatase activity. In this study, we present an *in vivo Drosophila* nephrocyte model of LS with a goal of understanding the sub-cellular mechanisms that may underlie renal dysfunction in these patients. Using a germline null allele of *dOCRL* (*dOCRL^KO^)* (Trivedi et al., 2020) and larval pericardial nephrocytes as a model, we find multiple defects in the endolysosomal system along with a physiological defect in clearing silver nitrate (AgNO_3_), a heavy metal toxin cleared via nephrocyte function. A nephrocyte specific deletion of *dOCRL*, demonstrated that these defects are cell autonomous to this cell type. The physiological defect in *dOCRL^KO^*nephrocytes was associated with multiple defects in the endosomal system including reduced endocytic cargo uptake, reduced early and late endosomal compartments as well as expanded but dysfunctional lysosomes. These sub-cellular defects as well as the physiological defect in AgNO_3_ clearance could be rescued by reconstituting *dOCRL^KO^* with dOCRL targeted selectively to the lysosomes. Thus, our studies define lysosomes as the primary site of function of dOCRL leading to overall homeostasis of the endosomal system in *Drosophila* nephrocytes.

## Results

### *dOCRL*^KO^ larvae show growth and developmental defects

In this study, we analysed *dOCRL^KO^*, using a previously generated protein null allele of *dOCRL* (Trivedi et al., 2020). Quantitative RT-PCR analysis showed virtually no *dOCRL* transcript in *dOCRL^KO^* animals **(Sup Fig 1A)** and a western blot with antibodies against dOCRL on larval lysates showed no detectable protein; protein levels could be restored by re-expression of HA::dOCRL **(Sup Fig 1C)**. Wandering larvae of *dOCRL^KO^*were smaller and weighed only about 50% of the body weight of controls **(Sup Fig 1B, 1F)**, showed a developmental delay of ca. 43 hours for 50% of animals to pupariation **(Sup Fig 1E)** and only about 58% of larvae pupariated **(Sup Fig 1G)**. *dOCRL^KO^* larvae showed lethality at every stage of development when grown on yeast medium **(Sup Fig 1D)**. The larval weight and larval lethality were rescued by the transgenic expression of a *dOCRL* using *hs-GAL4* that is expressed ubiquitously **(Sup Fig 1F, G)**. However, the adult lethality was not rescued, presumably due to insufficient expression of dOCRL with *hs-Gal4*. Lethality was however completely rescued (*data not shown*) by reconstitution using an X-duplication (BL-31454), which includes *dOCRL* genomic region. Collectively, these results suggest that *dOCRL* is required for normal growth and larval development in *Drosophila*.

**Figure 1:**
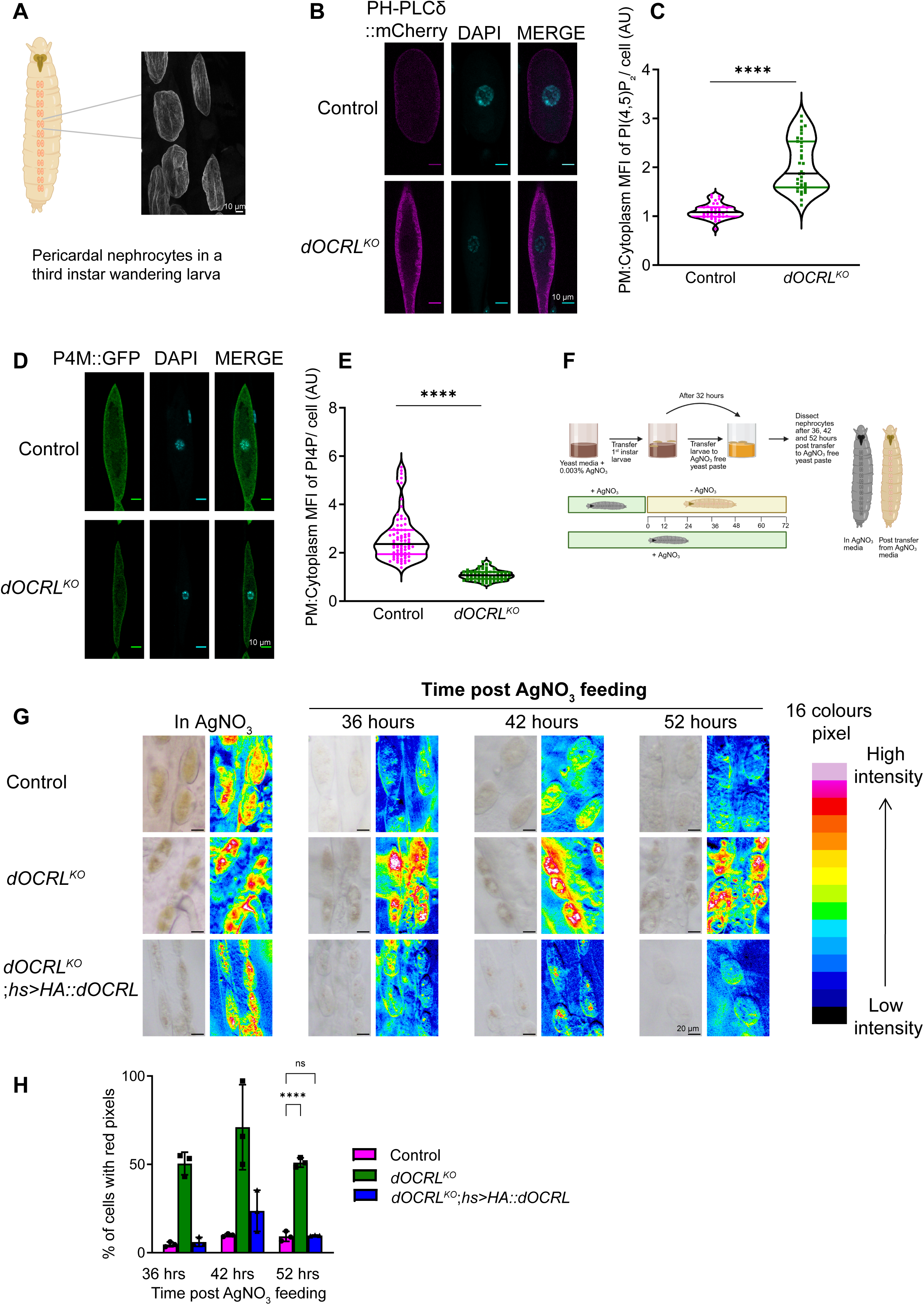
dOCRL regulates PI(4,5)P_2_ and PI4P levels and nephrocyte function. **(A)** Schematic showing the distribution of pericardial nephrocytes in a third instar larva. The micrograph shows a confocal section of nephrocytes labelled with antibodies to the slit diaphragm marker, pyd. Scale bar 10 μm **(B)** Confocal micrographs showing PI(4,5)P_2_ levels in control and *dOCRL^KO^*, detected with the PH-PLCδ::mCherry probe. Nucleus is marked by DAPI. Scale bar:10 μm. **(C)** Quantification of PI(4,5)P_2_ in control and *dOCRL^KO^* is shown. Y-axis shows the Mean Fluorescent Intensity (MFI) of plasma membrane to cytosolic ratio of PI(4,5)P_2_ in nephrocytes calculated using ImageJ. (*N*=9; *n*=27). Violin plots with mean and standard deviation are shown. **(D)** Confocal micrographs showing PI4P levels in control and *dOCRL^KO^* using P4M::GFP as a probe. Nucleus is marked by DAPI. Scale bar:10 μm. **(E)** Quantification of PI4P in control and *dOCRL^KO^* is shown. Y-axis shows the Mean Fluorescent Intensity (MFI) of plasma membrane to cytosolic ratio of PI4P in nephrocytes calculated using ImageJ. (*N* = 9; *n* = 68 ). Violin plots with mean and standard deviation are represented. **(F)** A cartoon illustrating the method of AgNO_3_ clearance assay. **(G)** Micrographs of nephrocytes from control, *dOCRL^KO^*, and *dOCRL^KO^;hs>HA::dOCRL* larvae showing AgNO_3_ uptake and cle*a*rance *a*t 36, 42, and 52 hours after transfer to AgNO_3_ free yeast paste. The corresponding 8-bit, 16-color pixel micrographs are displayed, with colours arranged by intensity from black (lowest) to white (highest). Scale bar: 20 μm. **(H)** Quantification of the percentage of cells displaying red pixel intensity, which corresponds to the highest levels of AgNO_3_ in nephrocytes for control, *dOCRL^KO^*, and *dOCRL^KO^;hs>HA::dOCRL*. The X-axis indicates time points at 36, 42, and 52 hours following transfer to AgNO_3_ free yeast paste, Y-axis shows the percentage of cells with red pixels. (N=9 biological replicates; each data point represents an individual trial; cells from all animals per genotype were pooled in each trial). Bars show mean and standard deviation. Statistical tests: (C and E) Student’s unpaired t-test is used. ****p<0.0001. (H) Two-way ANOVA grouped analysis with Dunnett’s multiple comparison tests performed using graph pad prism to compare between each group. ****p<0.0001, ***p<0.001, **p<0.01, ns-Non significance.

### *dOCRL* regulates plasma membrane PI(4,5)P₂ and PI4P in nephrocytes

OCRL is proposed to dephosphorylate PI(4,5)P_2_ to generate PI4P. We quantified the levels of PI(4,5)P_2_ at the plasma membrane of pericardial nephrocytes that are present around the heart tube **(Fig 1A)**. For this, we used the PH domain of PLCδ tagged to mCherry (PH-PLCδ::mCherry), a probe that specifically binds to PI(4,5)P_2_ (Hammond and Balla, 2015). We observed that in wild type nephrocytes, PH-PLCδ::mCherry uniformly decorates the plasma membrane **(Fig 1B)**. In *dOCRL^KO^* nephrocytes PH-PLCδ::mCherry accumulated at higher levels on the plasma membrane **(Fig 1B)**. The levels of PI(4,5)P_2_ at the plasma membrane were quantified by estimating the ratio of plasma membrane/cytoplasmic fluorescence and was higher in *dOCRL^KO^* compared to controls **(Fig 1C)**. Likewise, using the P4M::GFP probe that binds to PI4P (Balakrishnan et al., 2018) and mainly decorates the plasma membrane of nephrocytes **(Fig 1D)**, we estimated PI4P levels and found that PI4P level at the plasma membrane were lower in *dOCRL^KO^* compared to controls **(Fig 1E)**. Importantly, both the PH-PLCδ::mCherry and P4M::GFP probes were expressed equally in wild type and *dOCRL^KO^* **(Sup Fig 5 A-D)**.

### Depletion of *dOCRL* affects nephrocyte physiology

At the physiological level, nephrocytes maintain homeostasis by clearing out toxins from the haemolymph (Helmstädter et al., 2017). To investigate the contribution of dOCRL to the filtration function of nephrocytes, we measured the clearance of the heavy metal silver, using silver nitrate (AgNO_3_) from these cells, AgNO_3_ uptake has previously been used to study nephrocyte function (Fu et al., 2017a). For this, we transferred first instar larvae to feed on 0.003% AgNO_3_ in yeast paste and after 32 h, these larvae were transferred to yeast paste without AgNO_3_. Subsequently, larvae were removed from the AgNO_3_ free medium at defined time points, dissected and the amount of AgNO_3_ in each nephrocyte was visualized and quantified **(Fig 1F)**. This analysis was done on larvae at 36, 42, and 52 h post transfer from medium containing AgNO_3_ onto non-AgNO_3_ medium. Nephrocytes were imaged under a bright field microscope, and these images were converted to 8-bit images (16 color pixels) corresponding to the intensity of AgNO_3_, from white being highest concentration to black being least concentration. It was observed that in wild type animals, there were only traces of AgNO_3_ remaining by 36 h post-transfer and by 52 h AgNO_3_ was completely cleared out **(Fig 1G)**. However, in *dOCRL^KO^* nephrocytes, AgNO_3_ was not cleared at 36, 42, and 52 h post-transfer and a large proportion of nephrocytes containing AgNO_3_ could be observed; this clearance defect could be rescued by reconstituting with wild type *dOCRL* **(Fig 1G)**. We quantified the percentage of cells containing varying amounts of AgNO_3._ The micrographs were converted into 16 color pixels and the cells with high (red) levels of AgNO_3_ were counted. For each genotype we counted the proportion of cells with red pixels. The results, represented in (**Fig 1H)**, show that at all time points studied, *dOCRL^KO^* had a significantly higher percentage of cells with red pixels compared to wild type; this phenotype was rescued by reconstitution of *dOCRL^KO^* with wild type *dOCRL* **(Fig 1H)**. These findings demonstrate that *dOCRL* function is required for AgNO_3_ clearance in *Drosophila* nephrocytes.

### Rab7 depletion phenocopies the AgNO_3_ clearance defect in *dOCRL^KO^* nephrocytes

AgNO_3_ uptake is an established assay to investigate nephrocyte function in *Drosophila* (Kovalevskij, A, 1886). Although previous studies using this assay have addressed the role of Rab proteins in the endolysosomal system (Fu et al., 2017b), the precise mechanisms underlying cellular uptake and subsequent processing of AgNO_3_ in *Drosophila* pericardial nephrocytes remain to be elucidated. To determine the cellular basis of the AgNO_3_ clearance defect we have noted in *dOCRL^KO^*nephrocytes, we performed a genetic screen to test the potential role of dOCRL in nephrocytes. We curated a set of molecules that have previously been implicated in some aspect of the endolysosomal system and nephrocyte function **(Sup Table 1)**. We found that there was no AgNO_3_ uptake in nephrocytes when *shibire* (dynamin), a key regulator of clathrin mediated endocytosis was depleted **(Sup Table 1)**. Likewise, downregulating Rab5 function by expressing a dominant negative Rab5 (Rab5^DN^) also inhibited AgNO_3_ uptake **(Fig 2A)**. By contrast, expression of dominant negative Rab7 (Rab7^DN^) did not affect AgNO_3_ uptake but showed a clearance defect phenocopying *dOCRL^KO^* **(Fig 2 B,E)**. We screened several other candidates **(Sup Table 1)** with known roles in endolysosomal function but found no other molecule that could phenocopy *dOCRL^KO^*. In some cases, depleting or downregulating specific molecules led to grossly altered nephrocyte structure and function precluding any analysis of AgNO_3_ uptake and clearance.

**Figure 2:**
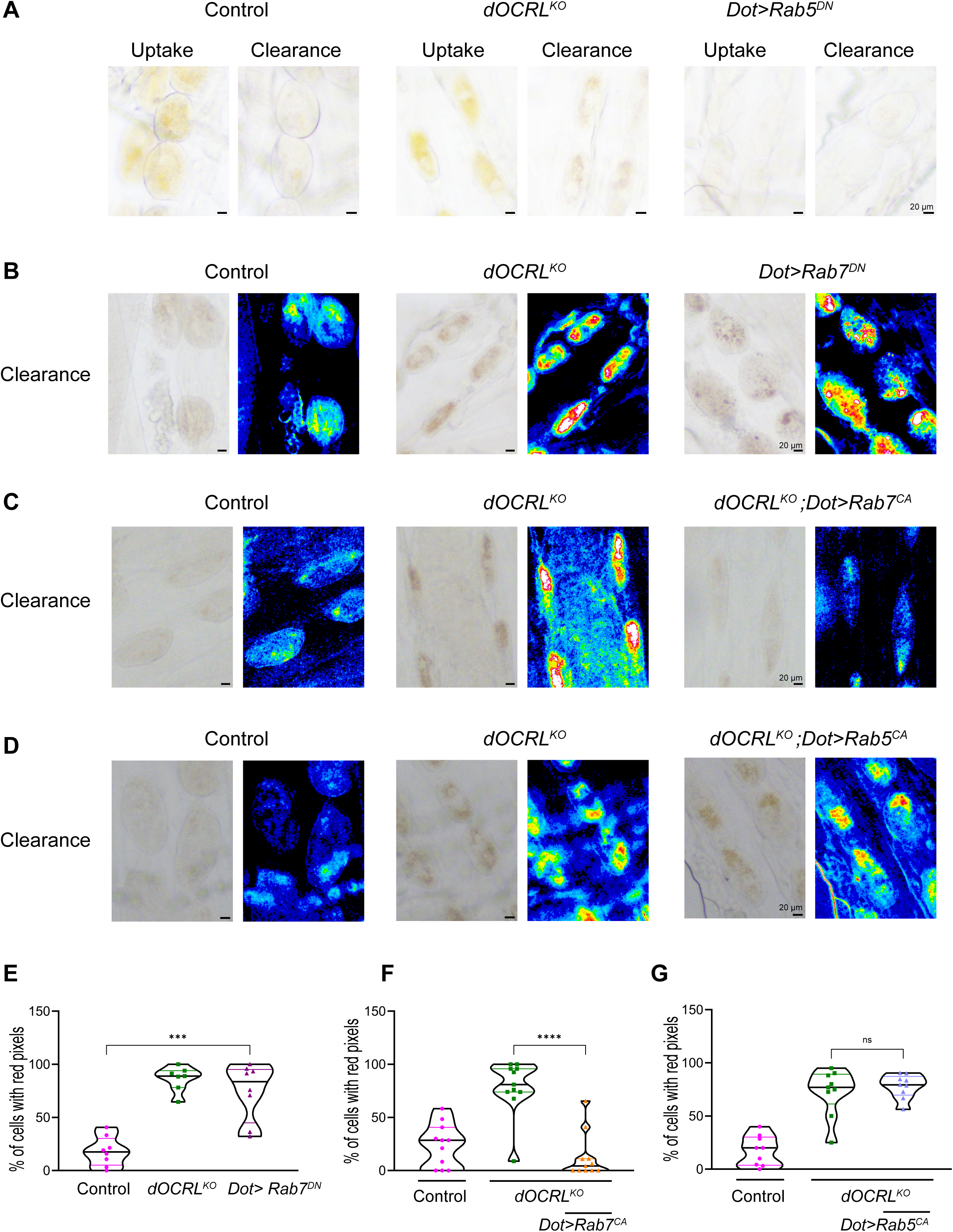
dOCRL is required at the late endolysosomal compartment to mediate AgNO_3_ clearance. **(A)** Micrographs of nephrocytes from control, *dOCRL^KO^* and *Dot>RAB5^DN^* larvae showing AgNO_3_ uptake, and subsequent clearance measured at 52 hours following transfer to AgNO_3_ free yeast media. (N=3; n=30). Scale bar:20 μm. **(B)** Micrographs of nephrocytes from control, *dOCRL^KO^* and *Dot>RAB7^DN^*larvae showing AgNO_3_ clearance at 52 hours after transfer to AgNO3 free yeast paste. The corresponding 8-bit, 16-color pixel micrographs are displayed. Scale bar: 20 μm. **(C)** Micrographs of nephrocytes from control, *dOCRL^KO^* and *dOCRL^KO^;Dot>RAB7^CA^*larvae showing AgNO_3_ clearance at 52 hours after transfer to AgNO_3_ free yeast paste. The corresponding 8-bit, 16-color pixel micrographs are displayed. Scale bar: 20 μm. **(D)** Micrographs of nephrocytes from control, *dOCRL^KO^*and *dOCRL^KO^; Dot>RAB5^CA^* larvae showing AgNO_3_ clearance at 52 hours after transfer to AgNO_3_ free yeast paste. The corresponding 8-bit, 16-color pixel micrographs are displayed. Scale bar: 20 μm. **(E)** Quantification of the percentage of cells exhibiting red pixel intensity, which corresponds to the highest levels of AgNO_3_ in nephrocytes for control, *dOCRL^KO^*, and *Dot>RAB7^DN^* at 52 hours after transfer. Each data point on the graph represents an individual animal, with 15–20 cells analysed per animal (N=8). Violin plots with mean and standard deviation are represented. **(F)** Quantification of the percentage of cells exhibiting red pixel intensity, which corresponds to the highest levels of AgNO_3_ in nephrocytes, is shown for control, *dOCRL^KO^*, and *dOCRL^KO^;Dot>RAB7^CA^* at 52 hours after transfer. Each data point on the graph represents an individual animal, with 15–20 cells analysed per animal (N=11). Violin plots with mean and standard deviation are represented. **(G)** Quantification of the percentage of cells exhibiting red pixel intensity, which corresponds to the highest levels of AgNO_3_ in nephrocytes, for control, *dOCRL^KO^*, and *dOCRL^KO^;Dot>RAB5 ^CA^* at 52 hours after transfer. Each data point on the graph represents an individual animal, with 15–20 cells analysed per animal (N=8). Violin plots with mean and standard deviation are represented. Statistical tests: (E-G) Students’s unpaired t-test is used. ns-Non significance, ****p<0.0001, ***p<0.001

To confirm the role of downregulated Rab7 function in the clearance defect of *dOCRL^KO^*, we expressed constitutively active Rab7 (Rab7^CA^) in *dOCRL^KO^* nephrocytes; this led to a rescue of the AgNO_3_ clearance defect in *dOCRL^KO^* **(Fig 2 C, F)**. By contrast, expression of constitutively active Rab5 could not rescue the clearance defect in *dOCRL^KO^* **(Fig 2 D, G)**. These findings suggest that reduced Rab7 function underlies the AgNO_3_ clearance defect in *dOCRL^KO^*.

### *dOCRL* is required for endocytosis at the plasma membrane

The endocytosis of fluid filtered through the slit diaphragm is a key part of nephrocyte function and it has been reported that nephrocytes internalize dextran by fluid phase endocytosis (Grawe et al., 2009). Since OCRL has been implicated in regulating endocytosis in cultured mammalian cells (Nández et al., 2014; Vicinanza et al., 2011), we studied this process using 10 kDa TMR-Dextran as a cargo in an *ex vivo* endocytosis assay. As a negative control, uptake assays were performed at 4^0^C since endocytosis is blocked at this temperature. At 25^0^C in wild type cells, robust endocytosis of TMR-Dextran was seen which was blocked at 4^0^C **(Fig 3 A, B)**. By contrast, in *dOCRL^KO^* there was almost complete absence of TMR-Dextran uptake at 25^0^C **(Fig 3 A, B)**. This defect in uptake could be rescued by reconstitution of *dOCRL^KO^* with *hs*>*HA::dOCRL* **(Fig 3A, B)**. We also assessed clathrin-mediated endocytosis in nephrocytes by using maleic anhydride-BSA (mBSA) conjugated to Cy5 (Haberland and Fogelman, 1985). In *dOCRL^KO^* nephrocytes, mBSA uptake was significantly reduced at 25^0^C compared to controls and was rescued when we reconstituted *dOCRL^KO^* with hs>*HA::dOCRL* **(Fig 3 C, D)**.

**Figure 3:**
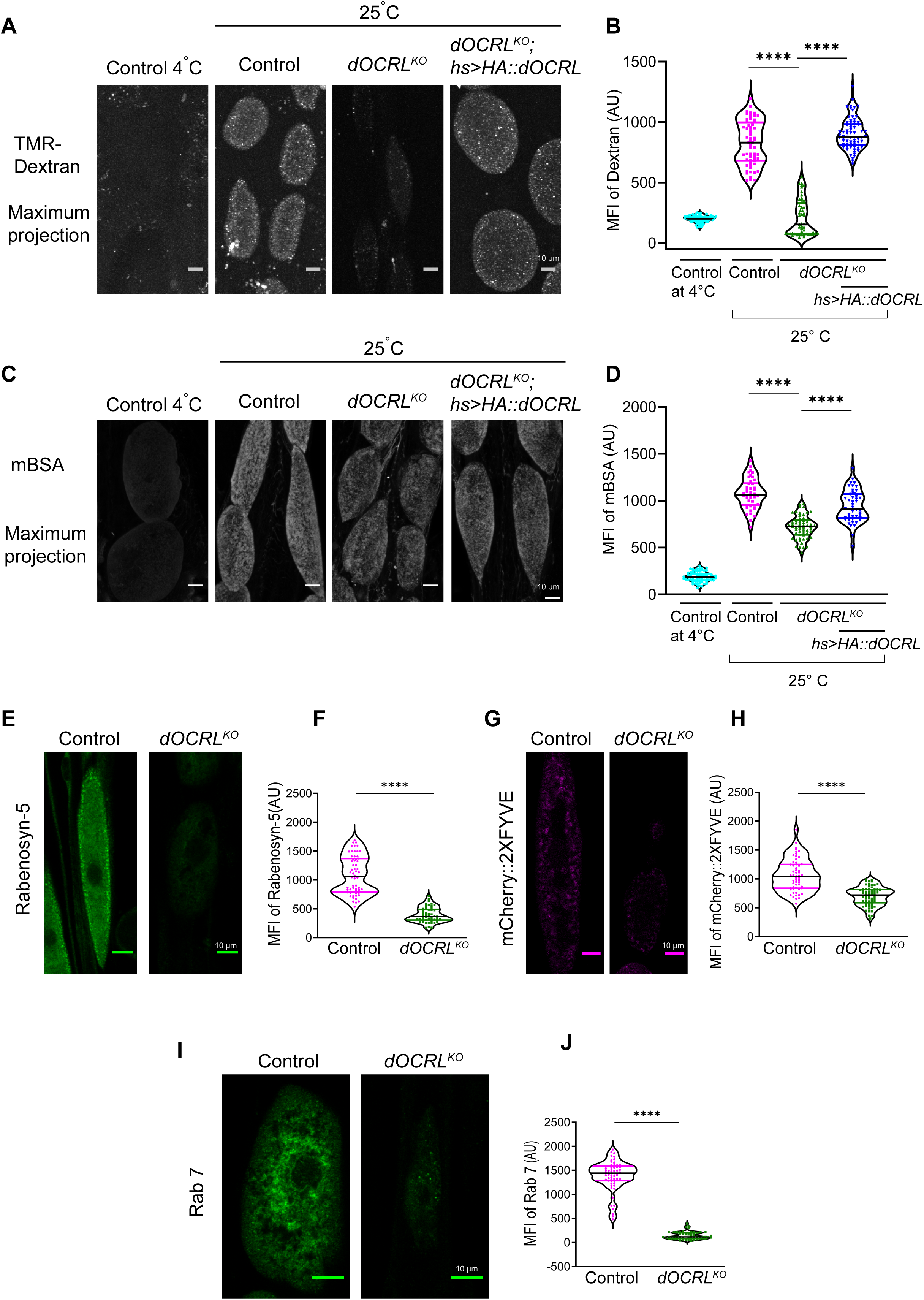
Loss of dOCRL perturbs endocytosis and endosomal homeostasis. **(A)** Confocal micrographs of 10kDa TMR-dextran uptake assay in control, *dOCRL^KO^*, *dOCRL^KO^;hs>HA::dOCRL.* Uptake at 4^0^C and 25^0^C is shown. Scale bar: 10 μm. **(B)** Quantification of dextran taken up by nephrocytes in control, *dOCRL^KO^* and *dOCRL^KO^;hs>HA::dOCRL*; Y-axis shows the Mean fluorescent intensity (MFI) of 10kDa TMR-Dextran in nephrocytes per unit area calculated using ImageJ (N=9) Each experiment consisted of three independent trials with three animals per group, and a total of 60 nephrocytes were imaged per trial. Violin plots with mean and standard deviation are represented. **(C)** Confocal micrographs of mBSA-Cy5 uptake assay in control, *dOCRL^KO^*, *dOCRL^KO^;hs>HA::dOCRL.* Scale bar: 10 μm. **(D)** Quantification of mBSA taken up by the cell in control, *dOCRL^KO^* and *dOCRL^KO^;hs>HA::dOCRL*; Y-axis shows the Mean fluorescent intensity (MFI) of mBSA-Cy5 in nephrocytes per unit area calculated using ImageJ (N=9) Each experiment consisted of three independent trials with three animals per group, and a total of 60 nephrocytes were imaged per trial. Violin plots with mean and standard deviation are represented. **(E)** Confocal micrographs showing immunostaining of control and *dOCRL^KO^* nephrocytes using the early endosome marker Rabenosyn-5. Scale bar: 10 μm. **(F)** Quantification of Rabenosyn-5 staining in control and *dOCRL^KO^*; Y-axis shows the Mean fluorescent intensity (MFI) of Rabenosyn-5 in nephrocytes per unit area calculated using ImageJ (N=9) Each experiment consists of three independent trials with three animals per group, and a total of 60 nephrocytes were imaged per trial. Violin plots with mean and standard deviation are represented. **(G)** Confocal micrographs of nephrocytes labeled with the mCherry::2XFYVE domain for control and *dOCRL^KO^*. Expression of UAS-mCherry::2XFYVE in nephrocytes was done using the Dot Gal4. Scale bar: 10 μm. **(H)** Quantification of mCherry::2XFYVE expression in control and *dOCRL^KO^*; Y-axis shows the Mean fluorescent intensity (MFI) of mCherry::2XFYVE in nephrocytes per unit area calculated using ImageJ (N=9). Each experiment consisted of three independent trials with three animals per group, and a total of 60 nephrocytes were imaged per trial. Violin plots with mean and standard deviation are represented. **(I)** Confocal micrographs showing immunostaining of control and *dOCRL^KO^* nephrocytes using antibodies to the late endosome marker Rab 7. Scale bar: 10 μm. **(J)** Quantification of Rab 7 staining in control and *dOCRL^KO^*; Y-axis shows the Mean fluorescent intensity (MFI) of Rab 7 in nephrocytes per unit area calculated using imageJ (N=9) Each experiment consisted of three independent trials with three animals per group, and a total of 60 nephrocytes were imaged per trial. Violin plots with mean and standard deviation are represented. Statistical tests: (B, D, F, H and J) Student’s unpaired t-test is used, ****p<0.0001.

Given the endocytic defects that we observed, we visualized endosomal compartments in *dOCRL^KO^* nephrocytes. Immunostaining with rabenosyn-5 (Mottola et al., 2010) a marker of early endosomes showed that rabenosyn-5 staining was substantially reduced in *dOCRL^KO^* compared to controls **(Fig 3E, F)**. The lipid phosphatidylinositol 3-phosphate (PI3P) is enriched on early endosomes and can be visualized using the 2XFYVE domain fused to a fluorescent protein (mCherry::2XFYVE); we found that the intensity of mCherry::2XFYVE punctae was lower in *dOCRL^KO^* compared to controls **(Fig 3G, H)**. The mCherry::2XFYVE probes were expressed equally in wild type and *dOCRL^KO^* (Sup Fig 5E, F). We also studied the late endosomal system in nephrocytes. Immunostaining for Rab7, a late endocytic marker, revealed that the total intensity of Rab7 staining, was lower in *dOCRL^KO^* **(Fig 3I, J)** suggesting that in nephrocytes, depletion of *dOCRL* impacts the late endocytic system. These results demonstrate that *dOCRL* is required for endosomal homeostasis.

### Altered late endosome-lysosome homeostasis in *dOCRL^KO^*

To visualize lysosomes, we used a reporter where the lysosome targeting sequence (LTS) is fused to mCherry (LTS::mCherry) (Ghosh et al., 2023). When expressed in control nephrocytes, LTS::mCherry marks punctate vesicular structures **(Fig 4A)** that can be quantified using confocal microscopy. We noted that the mean fluorescence intensity of LTS::mCherry was substantially elevated in *dOCRL^KO^* nephrocytes compared to control **(Fig 4A, B)**. The LTS::mCherry probes were expressed equally in wild type and *dOCRL^KO^* (Sup Fig 5G, H). We also found that the levels of cathepsin L, a lysosomal hydrolase was also upregulated in *dOCRL^KO^* **(Fig 4C, D)**. To test lysosomal function, we used the Magic Red™ stain which contains substrate of lysosomal proteases and gives fluorescence only when the substrate is cleaved by the lysosomal proteases, thus measuring lysosomal activity. We found a decrease in Magic Red™ fluorescence in *dOCRL^KO^* suggesting that lysosomal activity is reduced **(Fig 4E, F)**. Since lysosomal function is dependent on the pH of lysosomes, we stained the nephrocytes with lysotracker and found reduced fluorescence intensity in *dOCRL^KO^* **(Fig 4G, H)** suggesting a defect in the lysosomal acidification although the level of altered function is somewhat variable. To test if the reduced function of lysosomes also reflects in their reduced degradation of cargo, we conducted a pulse-chase analysis of mBSA localization with a longer chase duration of 40 minutes. We found increased mBSA intensity in *dOCRL^KO^*compared to control nephrocytes **(Fig 4I, J)** suggesting reduced degradation of this cargo in *dOCRL^KO^* nephrocytes.

**Figure 4:**
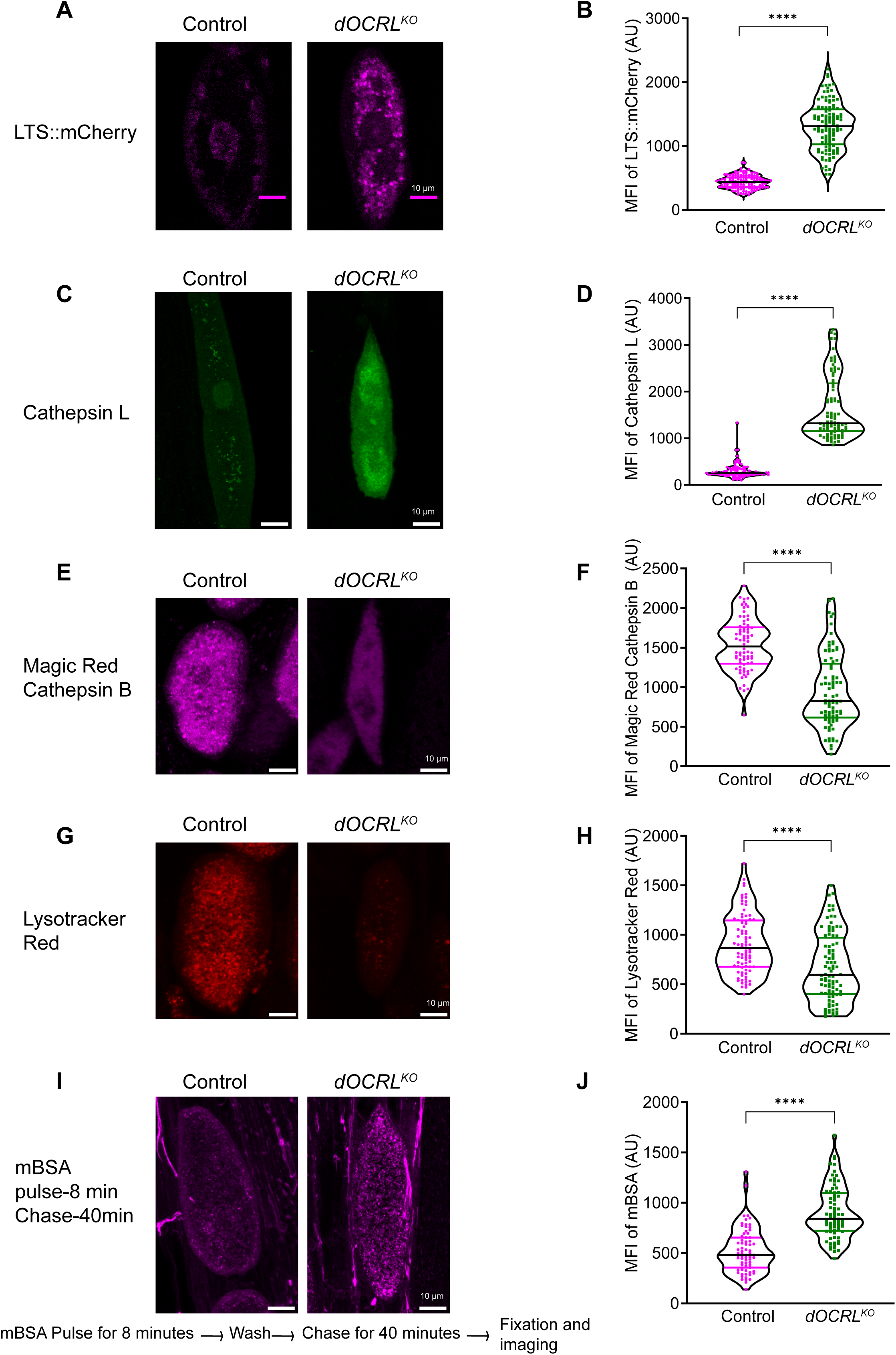
dOCRL is required to maintain lysosomal structure and function. **(A)** Confocal micrographs of nephrocytes labeled with the LTS::mCherry, are presented for control and *dOCRL^KO^*. Expression of UAS-LTS::mCherry in nephrocytes was done using Dot Gal4. Scale bar: 10 μm. **(B)** Quantification of LTS::mCherry expression in control and *dOCRL^KO^*; Y-axis shows the Mean fluorescent intensity (MFI) of LTS::mCherry in nephrocytes per unit area calculated using ImageJ (N=9, n=130). Violin plots with mean and standard deviation are represented. **(C)** Confocal micrographs showing immunostaining of control and *dOCRL^KO^* nephrocytes using the lysosomal marker Cathepsin L. Scale bar: 10 μm. **(D)** Quantification of Cathepsin L staining in control and *dOCRL^KO^*; Y-axis shows the Mean fluorescent intensity (MFI) of Cathepsin 7 in nephrocytes per unit area calculated using imageJ (N=9, n=90). Violin plots with mean and standard deviation are represented. **(E)** Confocal micrographs showing staining with Magic red cathepsin B substrate in control and *dOCRL^KO^* nephrocytes. Scale bar: 10 μm. **(F)** Quantification of Magic red cathepsin B substrate staining in control and *dOCRL^KO^*; Y-axis shows the Mean fluorescent intensity (MFI) of cathepsin B substrate in nephrocytes per unit area calculated using ImageJ (N=9, n=90). Violin plots with mean and standard deviation are represented. **(G)** Confocal micrographs showing staining of Lysotracker red in control and *dOCRL^KO^* nephrocytes. Scale bar: 10 μm. **(H)** Quantification of Lysotracker red staining in control and *dOCRL^KO^*; Y-axis shows the Mean fluorescent intensity (MFI) of Lysotracker red in nephrocytes per unit area calculated using ImageJ (N=9, n=90). Violin plots with mean and standard deviation are represented. **(I)** Confocal micrographs showing pulse-chase of mBSA in control and *dOCRL^KO^*nephrocytes. The *ex-vivo* nephrocyte preparation was pulsed with mBSA for 8 minutes followed by a chase duration of 40 minutes with the B & B buffer. Scale bar: 10 μm. **(J)** Quantification of mBSA in control and *dOCRL^KO^*; Y-axis shows the Mean fluorescent intensity (MFI) of mBSA in nephrocytes per unit area calculated using ImageJ (N=9, n=90). Violin plots with mean and standard deviation are represented. Statistical tests: (B, D, F, H, J) Student’s unpaired t-test is used. ****p<0.0001.

**Figure 5:**
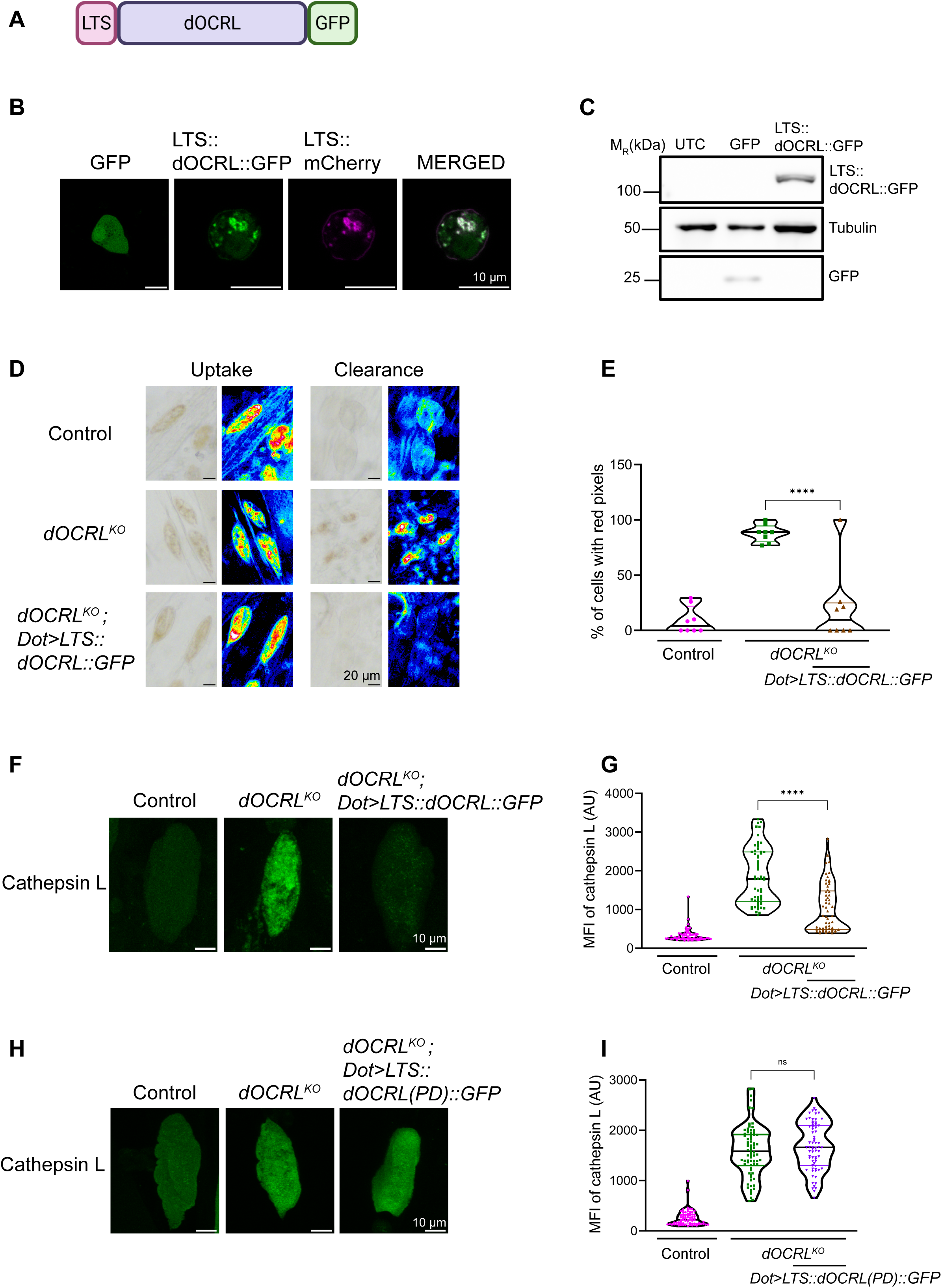
dOCRL localization to lysosomes in d*OCRL^KO^* is sufficient to facilitate AgNO_3_ clearance and restores lysosomal homeostasis. **(A)** Schematic representation of a construct designed to target dOCRL to the lysosomal compartment; LTS-lysosomal targeting signal (LTS); GFP Green Fluorescent Protein **(B)** Confocal micrographs of S2R+ cells expressing GFP alone, LTS::dOCRL::GFP and LTS::mCherry. **(C)** Immunoblots using S2R+cell lysates showing expression levels of LTS::dOCRL::GFP. The blot is probed with antibody against GFP and tubulin is used as a loading control. LTS::mCherry migrates at 130 kDa. (M_R_: molecular weight in kDa). **(D)** Micrographs of nephrocytes from control, *dOCRL^KO^* and *dOCRL^KO^; Dot>LTS::dOCRL::GFP* larvae showing AgNO_3_ clearance at 52 hours after transfer to AgNO_3_ free yeast paste. The corresponding 8-bit, 16-color pixel micrographs are displayed. Scale bar: 20 μm. **(E)** Quantification of the percentage of cells exhibiting red pixel intensity, which corresponds to the highest levels of AgNO_3_ in nephrocytes, is shown for control, *dOCRL^KO^* and *dOCRL^KO^; Dot>LTS::dOCRL::GFP* at 52 hours after transfer. Each data point on the graph represents an individual animal, with 15–20 cells analysed per animal (N=8). Violin plots with mean and standard deviation are represented. **(F)** Confocal micrographs showing immunostaining of control, *dOCRL^KO^* and *dOCRL^KO^; Dot>LTS::dOCRL::GFP* nephrocytes using the lysosomal marker Cathepsin L. Scale bar: 10 μm. **(G)** Quantification of Cathepsin L staining in control, *dOCRL^KO^* and *dOCRL^KO^; Dot>LTS::dOCRL*; Y-axis shows the Mean fluorescent intensity (MFI) of Cathepsin L in nephrocytes per unit area calculated using ImageJ (N=9, n=90). Violin plots with mean and standard deviation are represented. **(H)** Confocal micrographs showing immunostaining of control, *dOCRL^KO^* and *dOCRL^KO^; Dot>LTS::dOCRL(PD)::GFP* nephrocytes using the lysosomal marker Cathepsin L. Scale bar: 10 μm. **(I)** Quantification of Cathepsin L staining in control, *dOCRL^KO^* and *dOCRL^KO^; Dot>LTS::dOCRL(PD)::GFP*; Y-axis shows the Mean fluorescent intensity (MFI) of Cathepsin L in nephrocytes per unit area calculated using ImageJ (N=9, n=90). Violin plots with mean and standard deviation are represented. Statistical tests: (E, G, I) Student’s unpaired t-test is used. ****p<0.0001.

In the endolysosomal system, Rab7 positive endosomes fuse with lysosomes as a part of endosomal maturation. Given our finding of a reduced Rab7 compartment in *dOCRL^KO^* and AgNO_3_ clearance defects that were rescued by Rab7^CA^ expression, we tested whether expression of Rab7^CA^ in *dOCRL^KO^* could also rescue the lysosomal defect. To measure the lysosomal compartment, we measured the levels of cathepsin L. We found that the expression of Rab7^CA^ in the *dOCRL^KO^* was able to partially rescue the elevated levels of cathepsin L staining (**Sup Fig 3A, B**). This finding suggests that reduced Rab7 function likely contributes to the altered lysosomal defects and nephrocyte function in *dOCRL^KO^*.

### dOCRL at late endo-lysosomes rescues functional defects in *dOCRL^KO^* nephrocytes

The multiple compartments of the endolysosomal system are in a constant state of flux. Given our finding that loss of dOCRL leads to defects in early endosomes, late endosomes and lysosomes, we sought to determine the primary site where dOCRL function is required for endo-lysosomal homeostasis. For this we selectively targeted dOCRL to the late endosomal and lysosomal compartments. To do this, we designed a dOCRL transgene to target the protein selectively to lysosomes (by adding a lysosome targeting signal) (LTS::dOCRL::GFP) **(Fig 5A)**. The selective localization of the construct to lysosomes was verified by expressing it in *Drosophila* S2R+ cells. LTS::dOCRL::GFP was found to colocalize with the previous established probe for lysosomes LTS::mCherry (Ghosh et al., 2023) **(Fig 5B)**. Immunoblots with antibodies specific to GFP showed a band at 130 kDa for cells expressing LTS::dOCRL::GFP **(Fig 5C)**. We generated transgenic flies for expressing LTS::dOCRL::GFP. When expressed in nephrocytes, LTS::dOCRL::GFP was able to rescue the AgNO_3_ clearance defect **(Fig 5D, E)**. At the subcellular level, we found that LTS::dOCRL::GFP could partially rescue the elevated levels of cathepsin L staining in *dOCRL^KO^* **(Fig 5F, G)**. To determine whether dOCRL localized to lysosomes rescues the lysosomal accumulation defect through its phosphatase activity, specifically by regulating PI(4,5)P₂ levels, we conducted a multiple sequence alignment of *dOCRL* with human *OCRL* and *INPP5B* genes to identify conserved amino acid residues critical for enzymatic function **(Sup Fig 3C)**. The aspartic acid residue at position 523 in human OCRL is known to be essential for phosphatase activity (Nández et al., 2014). We identified and mutated the corresponding conserved aspartic acid at position 468 in dOCRL to glycine via site-directed mutagenesis. Transgenic flies expressing LTS::dOCRL(PD)::GFP were generated, the transgenic sequence was verified **(Sup Fig 3D)** and immunoblot analysis confirmed the expression of the phosphatase dead protein, both the LTS::dOCRL::GFP and LTS::dOCRL(PD)::GFP proteins were expressed equally in *dOCRL^KO^* **(Sup Fig 3D)**. The expression of LTS::dOCRL(PD)::GFP in *dOCRL^KO^* did not rescue the lysosomal accumulation defect **(Fig 5H, I)** assessed by Cathepsin L levels. This finding suggests that dOCRL restores lysosomal compartment upregulation in *dOCRL^KO^* through its catalytic phosphatase activity.

We also measured the size and function of the early endosomal compartment in *dOCRL^KO^* reconstituted with LTS::dOCRL::GFP. To determine the size of the early endosomal compartment, we performed Rabenosyn-5 staining and found that the reduced Rabenosyn-5 levels in *dOCRL^KO^* was restored back to wild type levels on reconstitution with LTS::dOCRL::GFP **(Fig Sup 4A, B)**. At a functional level, we studied the uptake of mBSA and found that the reduction in its uptake in *dOCRL^KO^* was also rescued on LTS::dOCRL::GFP expression **(Fig Sup 4C, D)**. Thus dOCRL function at the lysosome seems sufficient to rescue the defects at the early endosomal compartment seen in *dOCRL^KO^*.

### Cell autonomous regulation of nephrocyte function by dOCRL

*dOCRL^KO^* larvae show a whole-body growth phenotype along with defects in nephrocyte structure and function. To test if the requirement for *dOCRL* in supporting nephrocyte function is cell-autonomous, we generated a nephrocyte specific knockout of *dOCRL* using the CRISPR/Cas9 gene editing by selective expression of Cas9 (Trivedi et al., 2020); Cas9 was selectively expressed in nephrocytes using Dot GAL4. PCR analysis of larval carcasses of the appropriate genotype reveled a faint but clear *dOCRL* amplicon of 555 bp confirming the deletion of *dOCRL* **(Fig 6A)**; we refer to this allele as *dOCRL^N-KO^*. Using the PH-PLCδ::mCherry probe, we found that PI(4,5)P_2_ levels in *dOCRL^N-KO^* were significantly increased as compared to the control **(Fig 6C, D)**. We estimated nephrocyte function in *dOCRL^N-KO^* by measuring the fluid phase endocytosis in *dOCRL^N-KO^* and found reduced TMR-Dextran uptake phenocopying that seen in *dOCRL^KO^* **(Fig 6E, F)**. Dot GAL4, also shows some expression in salivary glands, lymph glands, and hemocytes. To exclude contributions from these non-nephrocyte tissues, we also generated *dOCRL* knockouts with AB1 GAL4 (salivary glands) and Hml GAL4 (hemocytes and lymph gland). No reduction in dextran uptake was observed in these experiments **(Sup Fig 2D-G)**. Additionally, a nephrocyte-specific knockout using Sns GAL4 confirmed that *dOCRL* loss reduces dextran uptake, consistent with results from Dot GAL4 **(Sup Fig 2H, I)**. Together, these results strongly suggest a cell-autonomous role for *dOCRL* dependent sub-cellular processes that underlie nephrocyte function.

**Figure 6:**
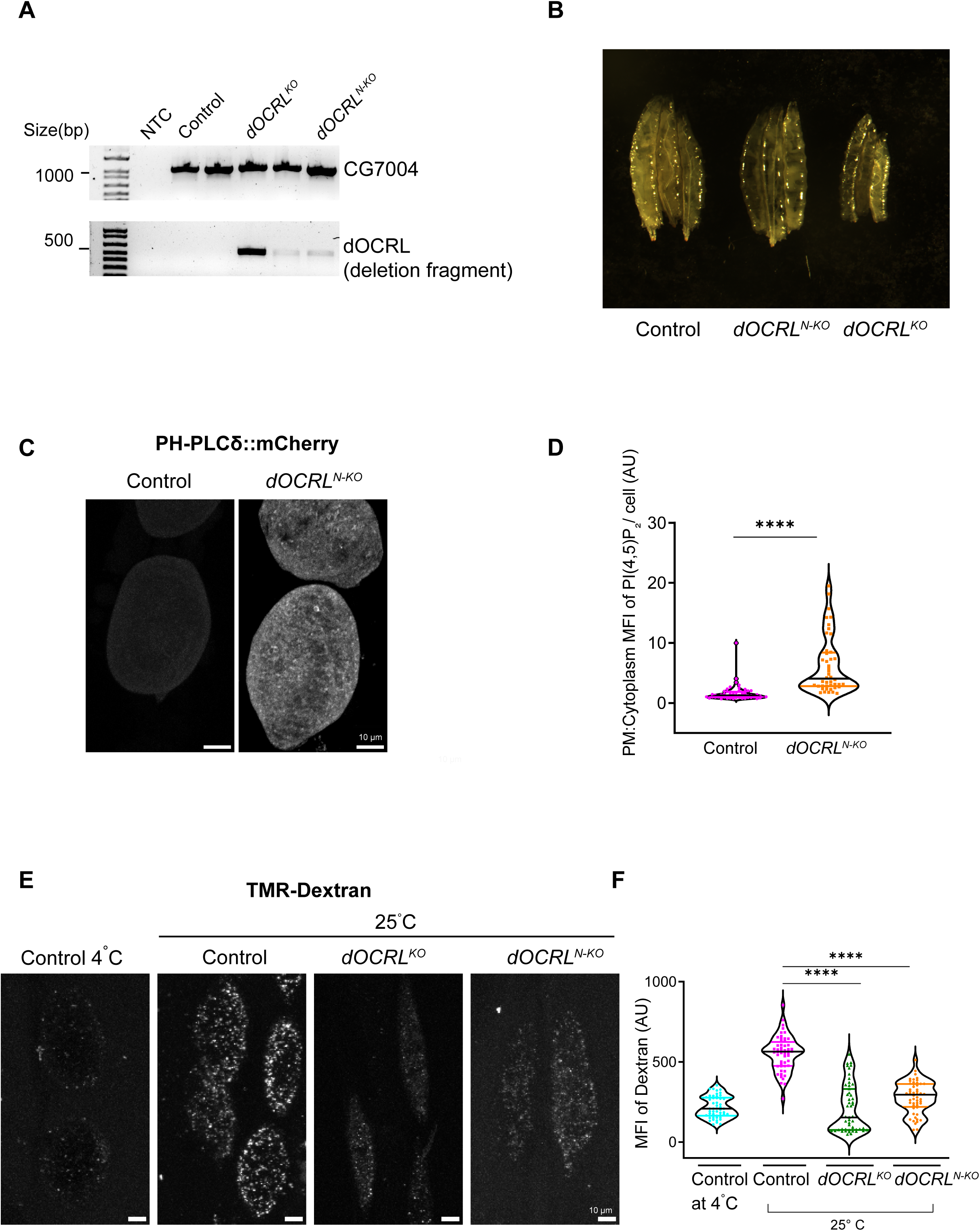
Generation of Nephrocyte specific deletion *dOCRL^N-KO^*. **(A)** Agarose gel electrophoresis to confirm the knockout of *dOCRL* gene in nephrocytes across three genotypes: control, *dOCRL^KO^* and *dOCRL^N-KO^*. The deletion fragment of the *dOCRL* gene was identified by its migration at 555 base pairs (bp) on the gel. Amplification of the CG7004 gene was included to serve as a control for DNA quality. **(B)** Micrographs of control, *dOCRL^N-KO^* and *dOCRL^KO^* third instar larvae. **(C)** Confocal micrographs showing PI(4,5)P_2_ levels in control, *dOCRL^N-KO^* detected with the PH-PLCδ::mCherry probe. Scale bar: 10 μm. **(D)** Quantification of PI(4,5)P_2_ in control and *dOCRL^N-KO^* is shown. Y-axis shows the Mean Fluorescent Intensity (MFI) of plasma membrane to cytosolic ratio of PI(4,5)P_2_ in nephrocytes calculated using ImageJ. (*N* = 9; *n* = 50). Violin plots with mean and standard deviation is shown. **(E)** Confocal micrographs of 10kDa TMR-dextran uptake in control, *dOCRL^KO^* and *dOCRL^N-KO^.* Scale bar: 10 μm. **(F)** Quantification of dextran taken up by the cell in control, *dOCRL^KO^*and *dOCRL^N-KO^*; Y-axis shows the Mean fluorescent intensity (MFI) of 10kDa TMR-Dextran in nephrocytes per unit area calculated using ImageJ (N=9). Each experiment consisted of three independent trials with three animals per group, and a total of 50 nephrocytes were imaged per trial. Violin plots with mean and standard deviation are represented. Statistical tests: (D, F) Student’s unpaired t-test is used. ****p<0.0001.

We also monitored growth and development in *dOCRL^N-KO^*animals. The weight and size of *dOCRL^N-KO^* 3^rd^ instar larvae was only modestly lower than that of controls **(Fig 6B, Sup Fig 2 C)**. *dOCRL^N-KO^* larvae completed larval development, underwent pupal metamorphosis **(Sup Fig 2A)** and eclosed as adults. However, only a proportion of flies eclosed, and the remaining did not complete pupal metamorphosis **(Sup Fig 2 B)**.

## Discussion

In LS, the function of PTC of the kidney is known to be impaired. The function of the endosomal system is critical to the function of the PTC. Following glomerular filtration, the uptake of fluid and proteins by PTC is critical for normal renal function and this uptake is dependent on endocytosis. Several studies have reported that in OCRL depleted cells, the process of endocytosis is impaired. The endosomal system is connected to lysosomes that have also been shown to be abnormal in OCRL depleted cells. The OCRL protein has been localized at both early endosomes and lysosomes, and this raises a question of which of these locations it functions at to support PTC function. In this study, we have used *Drosophila* nephrocytes as a model to investigate this question. Depletion of OCRL in pericardial nephrocytes recapitulates phenotypes of OCRL depletion in human cells at multiple scales of analysis- (i) elevation of plasma membrane PI(4,5)P_2_ and a reduction of PI4P levels (ii) Alteration in the size of the early endosomal compartment along with reduced endocytic uptake (iii) increase in the size of the lysosomal compartment along with lysosome dysfunction. Therefore, *Drosophila* nephrocytes are a suitable model to perform a multiscale analysis of the impact of OCRL loss on PTC function.

How does loss of OCRL function impact nephrocyte function? One possibility is that the uptake of AgNO_3_ from hemolymph by endocytosis is impacted by OCRL depletion; however, in this study we found that AgNO_3_ uptake was not affected in *dOCRL^KO^*; this contrasts with downregulation of Rab5 function which led to a dramatic reduction in AgNO_3_ uptake into nephrocytes **(Sup Table 1)**. Thus, the reduced endocytic uptake noted in *dOCRL^KO^* nephrocytes is unlikely to underlie their impaired function. An equivalent study in mice has not been reported.

Rab GTPases are key regulators of the endolysosomal system. To elucidate where in the endosomal system of nephrocytes OCRL function might be critical, we performed a systematic screen of Rab proteins and found that downregulation of Rab7 in wild type nephrocytes phenocopies the effects of OCRL depletion on AgNO_3_ handling; i.e no detectable effect on AgNO_3_ uptake but a defect in its clearance from nephrocytes. Rab7 is a GTPase that regulates multiple functions including early-late endosome maturation, late endosome-lysosome fusion and autophagosome-lysosome fusion (Guerra and Bucci, 2016). In *dOCRL^KO^* nephrocytes, we noted that the early and late endosomal compartment were not expanded; these findings suggest that it is unlikely that maturation from early to late endosomes is regulated by OCRL. Rather, we found that the lysosomal compartment was expanded and dysfunctional. As OCRL has been localized on the lysosomal membrane, it is possible that the primary defect in *dOCRL^KO^*is altered lysosomal function. However, since the endo-lysosomal system is inter-connected and in a state of homeostatic flux, it is possible that the primary defect in OCRL depleted nephrocytes is altered lysosomal function and the altered early endosome compartment and function is a compensatory effect on the endosomal system. Alternatively, it is possible that in nephrocytes, OCRL functions at both the early endosomal compartment and in lysosomes. We found that reconstituting *dOCRL^KO^* with a dOCRL protein targeted exclusively to the lysosomal compartment led to a rescue of the lysosomal defects underscoring a primary role for dOCRL function at this organelle membrane. Remarkably, this reconstitution of *dOCRL^KO^* with lysosome targeted dOCRL also resulted in a rescue of the endocytic uptake defect and early endosome abnormalities. Conversely, expression of dOCRL selectively targeted to early endosomes did not rescue the early endosomal defects in *dOCRL^KO^* (data not shown). Taken together, these findings strongly suggest that the primary defect in the endolysosomal system of *dOCRL^KO^* is the lack of dOCRL enzymatic activity at the lysosome and the endocytic defects are secondary to this primary dysfunction.

In summary, this study presents a multiscale analysis of the biology of LS in a *Drosophila* nephrocyte model. We find that that dOCRL is required to regulate lysosomal function and endomembrane homeostasis. This in turn is required to deliver normal nephrocyte function. Our work opens the way forward to both mechanistic studies as well as screening for new compounds that could correct this defect in lysosomal function and therefore be developed into new therapeutic agents for managing the renal defects in LS.

## Materials and methods

### *Drosophila* culture and strains

All flies used in this study were reared at 25^0^ C and 50% relative humidity on standard cornmeal media (Per litre of media contains 80 g corn flour, 20 g D-Glucose, 40 g Sucrose, 15 g yeast powder, 8 g agar, 4 ml propionic acid, 0.6 ml orthophosphoric acid and 1 g Methyl parahydroxy benzoate) in a laboratory incubator. All experiments were performed with third instar wandering larvae grown on yeast paste layered on 1% bacteriological agar. The following transgenic lines were obtained from Bloomington *Drosophila* stock center: Red Oregon-R (ROR)-wild type control (BL-2376), *Dot-GAL4* (BL-6903), *Dot-GAL4* (BL-67609), X-duplication (BL-31454), *UAS-Fab1 RNAi* (BL-35793), *UAS-Rab2-DN* (BL-23640), *UAS-Rab5-DN* (BL-9772), *UAS-Rab7-DN* (BL-9778), *UAS-Rab7-CA* (BL-9779), *UAS-Rab11-DN* (BL-23261), *UAS-Rab4-DN* (BL-9768), *UAS-Rab27-DN* (BL-23267), *UAS-Shibire* RNAi (BL-28513), *UAS-Trpml* RNAi (BL-31294), *Trpml^KO^* (BL-28992), *UAS-VHA-*39 RNAi (BL-35029), *UAS-syntaxin17* RNAi (BL-25896), *SNS-GAL4* (BL-92192), *AB1-GAL4* (BL-1284), *HML-GAL4* (BL-30140), *UAS-Clc-c/Clcn5* RNAi (BL-27034), *Clc-b/Clcn7^KO^* (BL-80059), *CP1/CathepsinL^KO^*(BL-51555), *UAS-*ATG1 overexpression (BL-51655). Additionally, the following transgenic lines were used: *UAS-mCherry::2XFYVE* (Amy Kiger, UCSD), *UAS-PH-PLCδ::mCherry* (Patrick Verstreken lab), *UAS-P4M::GFP* (Balakrishnan et al., 2018) and *UAS-LTS::mCherry* (Ghosh et al., 2023). For the generation of nephrocyte-specific *dOCRL* knockout (*dOCRL^N-KO^*) flies, lines described in Trivedi et al., 2020, *gRNAP2dual-CG3573V* (BDSC_92492) and *UAS-Cas9-T2A-eGFP* (BDSC_94298) were used.

### Generation of transgenic flies

The transgenic strains *HA::dOCRL* were generated by cloning *dOCRL* cDNA (RRID:DGRC_1325630) into pUAST-attB (RRID:DGRC_1419) with HA tag. To generate the lysosomal targeted construct of dOCRL, a stepwise cloning strategy was employed. Initially, a construct *LTS::GFP* was generated using Gibson assembly. In this process, the LTS fragment was amplified utilizing *Lyso-dPIP4K::eGFP* (Sharma et al., 2019) as the template. The amplified LTS fragment was then ligated into the pUAST-attB vector, containing *XhoI* and *NotI* restriction sites using Gibson assembly. Following the generation of the *LTS::GFP* construct, restriction digestion was performed using *XhoI*, allowing the construct to be used as a vector for further cloning. The dOCRL fragment was amplified from the pTHW-dOCRL construct (lab generated). Using Gibson assembly, the amplified *dOCRL* fragment was ligated into the digested *LTS::GFP* vector, resulting in the generation of the final construct, *LTS::dOCRL::GFP*. Transgenic lines were generated using site-specific recombination into attP sites.

### Site directed mutagenesis

To generate the *LTS::dOCRL(PD)::GFP* transgene, mutations were introduced into the *LTS::dOCRL::GFP* containing pUAST plasmid. The conserved amino acid required for enzymatic activity in dOCRL was identified via multiple sequence alignment (by Clustal Omega (RRID:SCR_001591) with the human *OCRL* and *INPP5B* gene. Amino acid 523 in hOCRL is reported to contribute to phosphatase activity in human OCRL (Nández et al., 2014). A corresponding mutation was made at amino acid 468 in dOCRL, which contains the same conserved residue, by site-directed mutagenesis. Aspartic acid residue was substituted with glycine to abolish phosphatase activity in dOCRL. Mutations were incorporated into the primer pair, and PCR-based site-directed mutagenesis (SDM) was carried out to introduce point mutations into the *LTS::dOCRL::GFP* construct. During the first 10 cycles of PCR, the forward or reverse primers containing the mutations were used in two separate reactions; these reactions were then combined for the remaining 15 cycles to complete the 25-cycle protocol.

The *PCR* product was digested with *DpnI* to remove the parent vector. The resulting DNA was transformed, and colonies were inoculated into LB media for overnight incubation at 37°C with shaking. Plasmid isolation was performed the next day, and samples were submitted for Sanger sequencing to confirm the presence of the mutation. The primers used to introduce the PD mutation are as follows :-

dOCRL-D-G-FP GAGATTCGGCAGAGCGGTCACAAGCCCGTGTA

dOCRL-D-G-RP TACACGGGCTTGTGACCGCTCTGCCGAATCTC

### S2R+ cell culture

*Drosophila* S2R+ (gift from Jitu Mayor lab) cells were transfected with GFP or mCherry-tagged constructs using Effectene (Qiagen #301425). Transfected cells were cultured for 48 hours prior to being plated in dishes for confocal imaging or for protein isolation to perform immunoblotting. For imaging, cells seeded onto coverslip dishes were allowed to settle for one hour. The cells were then fixed with 2.5% paraformaldehyde (PFA) (EMS #15710) at room temperature for 20 minutes, followed by washes with phosphate buffered saline (PBS). Imaging of S2R+ cells was conducted using an FV3000 confocal microscope (RRID:SCR_017015) at 60X magnification.

### Quantitative RT-PCR

Total RNA was extracted from third instar wandering larvae using TRIzol reagent (Ambion, #15596018) followed by treatment with DNase-I (Invitrogen, #18068015). cDNA was synthesized using Superscript II RNase H-Reverse transcriptase (Invitrogen, #18064014) and random hexamers (Invitrogen, #N8080127). Both non-template controls and no reverse transcription controls were included in the analysis. Quantitative PCR was performed using cDNA samples and primers using Power SYBR Green PCR Master Mix (Applied biosystems, #4367659) on an Applied Biosystem 7500 Fast Real-Time PCR system (RRID:SCR_018051). Expression levels were normalized to the Ribosomal Protein 49 (*RP49*), a housekeeping gene. Primers were designed at exon-exon junction and met all recommended qPCR design parameters.

The primers used for q-PCR were: RP49 forward: CGGATCGATATGCTAAGCTGT; RP49 reverse: GCGCTTGTTCGATCCGTA; dOCRL forward: GAACAACAAGACCTGCAGC; dOCRL Reverse: CTGTCCATCATCTTATCGATCC

### Growth curve assay

In this assay, both control and *dOCRL^KO^* flies were permitted to lay eggs for 4–6 hours on standard fly food. At 24 hours after egg laying, first instar larvae were transferred to yeast media under controlled crowding conditions, with approximately 25 larvae placed per vial. Six biological replicates were maintained and incubated at 25^0^C. After 96 hours, the number of larvae that had pupariated was recorded for both groups at three-hour intervals. The proportion of larvae that pupariated was calculated and used to determine the pupariation percentage for each time bin. These percentages were then plotted and fitted using a variable slope model in GraphPad Prism (RRID:SCR_002798).

### Nephrocyte function assay

In this assay, flies were allowed to lay eggs on standard fly food. After 24 hours, first instar larvae were transferred to yeast paste containing either 0.0015% (in figure 2 and 5 D) or 0.003% (in figure 1 G) AgNO_3_ (Sigma, #209139), layered on 1% bacteriological agar. Following 32 hours of feeding on AgNO_3_ media, larvae were transferred to yeast paste media without AgNO_3_. At 36, 42, and 52 hours after transfer to the AgNO_3_ free yeast paste, nephrocytes were dissected, fixed with 4% paraformaldehyde for 15 minutes at room temperature, washed three times with PBS, and mounted in 70% glycerol. Images were captured with a 10X objective on an Olympus BX53 microscope (RRID:SCR_022568) using Olympus cellSens Software (RRID:SCR_014551). These images were converted to 8-bit, 16-color pixel images corresponding to the intensity of AgNO_3_, from white (highest) to black (lowest). Cells with red pixels (high AgNO_3_) were counted and results were plotted using Graphpad Prism (RRID:SCR_002798).

### mBSA uptake assay

mBSA was prepared as described earlier (Haberland and Fogelman, 1985) and conjugated with Cy5 bis-functional dye (GeneCopoeia, #C183) according to manufacturer’s instructions. Pericardial nephrocytes in larval fillet were dissected in Brodie and Bate’s buffer (B & B buffer) (135 mM NaCl, 5 mM KCl, 4mM MgCl_2_, 2mM CaCl_2_, 5mM TES, 36mM sucrose) and were incubated with conjugated mBSA (0.1mg/ml) for 15 minutes at 25^0^C in dark. Subsequently, cells were washed with PBS, fixed with 4% PFA for 15 minutes at room temperature. The tissues were washed with PBS thrice and mounted onto glass slides using 70% glycerol in PBS. For mBSA pulse-chase assay, the nephrocytes were pulsed with mBSA in B & B buffer for 8 minutes. This was followed by washing and a chase period of 40 minutes in just B & B buffer at 25^0^C in the dark, followed by fixation and imaging. Imaging was performed using an Olympus FV3000 confocal microscope (RRID:SCR_017015). ImageJ was used for quantification of mean fluorescence mBSA intensity per unit area of nephrocytes from maximum projection images.

### Dextran uptake assay

Pericardial nephrocytes in larval fillet were dissected in B & B buffer. The samples were incubated in 0.33 mg/ml of TMR-dextran (Thermo-fisher scientific, # D1868) for 5 minutes in B & B buffer at 25^0^C in dark. For the negative control, a separate set was incubated with dextran at 4^0^C. Following incubation, samples were washed in ice-cold PBS for 10 minutes, fixed in 4% paraformaldehyde (PFA) in PBS for 15 minutes at room temperature (25^0^C), and subsequently washed three times with PBS. The tissues were then mounted onto glass slides using 70% glycerol in PBS. Imaging was performed using an Olympus FV3000 confocal microscope (RRID:SCR_017015). ImageJ was used for quantification of mean fluorescence dextran intensity per unit area of nephrocytes from maximum projection images.

### Western Blot

All immunoblots used lysates from third instar wandering larvae, except for probe level estimation, where larval fillets with nephrocytes were dissected in PBS and processed for protein extraction. The samples were homogenized in Tris-Lysis buffer containing Protease inhibitor (Roche, #04693142001) and PhosStop (Roche, #04906845001) and heated at 95^°^C for 5 minutes. Protein extracts were separated by SDS-Polyacrylamide gel electrophoresis and transferred onto nitrocellulose filter membrane (Hybond-C Extra; [GE Healthcare]) using a wet transfer apparatus (BioRad). Membranes were blocked with 5% Blotto (Santa Cruz Biotechnology, #sc-2325), in phosphate buffer saline (PBS) with 0.1% Tween 20 (Sigma Aldrich, #P1379) (PBST) for 2 h at room temperature (25^0^C) followed by incubation in primary antibody diluted in 5% Blotto in PBST incubated overnight at 4^0^C. Primary antibodies dilutions used were 1:1000 anti-dOCRL (Avital Rodal lab), 1:4000 anti-beta-tubulin (DHSB, #E7c, RRID: AB_528499), 1:1000 anti-Syntaxin1A (DSHB, #8c3 RRID: AB_528484), 1:1000 anti-mcherry (Thermo-fisher scientific, #PA5-34974, RRID: AB_2552323), 1:2000 anti-GFP (Santa Cruz Biotechnology, #sc9996, RRID: AB_627695). Following this, the membrane was washed for 10 minutes thrice with PBST and incubated at 1:10000 dilution with the appropriate HRP conjugated secondary antibody: Peroxidase Affinipure Goat Anti-Mouse IgG (H+L) (Jackson ImmunoResearch Laboratories, #115-035-003, RRID: AB_10015289), Peroxidase Affinipure Donkey Anti-Rabbit IgG (H+L) (Jackson ImmunoResearch Laboratories, #711-035-152, RRID: AB_10015282) in PBST for 2 hours at room temperature. Following this, the membrane was washed for 10 minutes thrice with PBST. Blots were developed with ECL substrate (Bio-Rad, #1705061) and imaged using LAS 4000 ImageQuant (RRID:SCR_014246) (GE Healthcare). Whenever required, blots were stripped with 3% glacial acetic acid in PBST for 30 minutes, washed three times, and re-probed with primary antibody.

### Immunostaining of nephrocytes

Pericardial nephrocytes in larval fillet were dissected from wandering third instar larvae in B & B buffer and fixed in 4% paraformaldehyde in PBS for 15 minutes at room temperature. After fixation, cells were rinsed thrice with 0.3% PBTX (0.3% Triton X in PBS) for 10 minutes. The samples were incubated in blocking solution (5% Fetal Bovine Serum in PBTX) for two hours at room temperature. They were then incubated overnight at 4^0^C with the appropriate primary antibody: anti-Rab7, 1:200 (DSHB, RRID: AB_2722471); anti-Rabenosyn-5, 1:800 [gift from Marino Zerial, MPI-CBG, Dresden, Germany; (Mottola et al., 2010)]; and anti-cathepsin L (Abcam, #ab58991, RRID: AB_940826). The samples were washed thrice with 0.3% PBTX and were subsequently incubated at room temperature for four hours with the appropriate secondary antibodies: Goat anti-Mouse IgG [H+L] Cross-Adsorbed Secondary Antibody, Alexa Fluor 488 (Thermo Fisher Scientific, Cat# A-11001, RRID: AB_2534069), Goat anti-Rabbit IgG [H+L] Highly Cross-Adsorbed Secondary Antibody, Alexa Fluor 488 (Thermo Fisher Scientific, Cat# A-11034, RRID: AB_2576217), Donkey anti-Rabbit IgG [H+L] Highly Cross-Adsorbed Secondary Antibody, Alexa Fluor 568 (Thermo Fisher Scientific, Cat# A10042, RRID: AB_2534017) at 1:300 dilution. Samples were washed three times in 0.3% PBTX, rinsed once in PBS, and mounted in 70% glycerol in 1X PBS. Imaging was performed using an Olympus FV3000 confocal microscope (RRID:SCR_017015). ImageJ was used for quantification of mean fluorescence intensity per unit area of nephrocytes from maximum projection images.

### Lysotracker Assay

Larval fillet preparations for nephrocytes were performed using B & B buffer, followed by incubation with Lysotracker Red (Thermo Fisher Scientific, #L7528) at a 1:1000 dilution in B & B buffer for 15 minutes at room temperature. Samples were then washed three times in ice-cold PBS, fixed with 1% paraformaldehyde in PBS for 2 minutes, and mounted on glass slides with 70% glycerol in PBS. Imaging was conducted immediately using an Olympus FV3000 confocal microscope (RRID:SCR_017015), and mean fluorescence intensity per unit area of nephrocytes was quantified with Fiji ImageJ (RRID:SCR_003070).

### Magic red cathepsin B assay

Larval fillet preparations for nephrocytes were performed using B & B buffer, followed by incubation with Magic Red stain (Bio-Rad, #ICT937) at a 1:25 final dilution in B & B buffer for 30 minutes in the dark at 25^0^C. Samples were then washed and mounted in B & B buffer onto glass slides. Imaging was conducted immediately using an Olympus FV3000 confocal microscope (RRID:SCR_017015), and mean fluorescence intensity per unit area of nephrocytes was quantified with Fiji ImageJ (RRID:SCR_003070).

### DNA extraction for verification of *dOCRL^N-KO^*

For the verification of *dOCRL^N-KO^*, eight control and *dOCRL^N-KO^* third instar wandering larvae were dissected to obtain larval fillets containing only nephrocytes, prepared in B & B buffer. Following the removal of the buffer from the dissected tissues, the carcasses were transferred into a 0.5 ml microcentrifuge tube. The collected samples were homogenized in 50 µl of squishing buffer supplemented with 200 µg/ml Proteinase K (Sigma, #P2308) using a pipette tip. The homogenates were incubated at room temperature (25^0^C) for 30 minutes. Subsequently, Proteinase K was inactivated by heating the samples at 95°C for 5 minutes. 3µl of this lysate was used as template for verification of *dOCRL^N-KO^*by polymerase chain reaction analysis. We used CG7004 gene as a control to determine the quality of DNA sample preparation.

### Statistical Analysis

Each experiment was conducted at least twice with multiple biological replicates. Statistical analyses were performed using GraphPad Prism (RRID:SCR_002798). For the comparison between two experimental groups, Student’s unpaired t-test was employed. When experiments involved the comparison of more than two groups, a two-way ANOVA grouped analysis followed by Dunnett’s or Tukey’s multiple comparison tests was performed to determine statistical significance. For survival analysis, the Long-rank (Mantel-Cox) test and the Gehan-Breslow-Wilcoxon test were utilized to assess differences in survival distributions between two genotypes.

## Acknowledgements

This study was supported by the Department of Atomic Energy, Government of India (RTI-0046) and a Department of Biotechnology, Government of India grant to AV (BT/PR11030/MED/30/1644/2016). We thank the *Drosophila* facility, Imaging Facility and Sequencing facility at NCBS-TIFR for support.

## Supplementary Figures

**Supplementary figure 1: Growth and development phenotypes of *dOCRL^KO^*larvae (A)** Quantitative real-time PCR analysis of *dOCRL* transcript levels in control and *dOCRL^KO^.* Y-axis indicates dOCRL transcript level expression normalized to the loading control RP49 (Ribosomal protein 49), n=4 biological replicates. **(B)** Micrographs of control, *dOCRL^KO^*, and *dOCRL^KO^;hs>HA::dOCRL* third instar larvae. **(C)** Immunoblots from third instar wandering larvae of genotypes indicated at the top of the blot. The blot was probed with an antibody against dOCRL and tubulin was used as a loading control. dOCRL migrates at 100 kDa. (M_R_: molecular weight in kDa). **(D)** Survival plot of control and *dOCRL^KO^* larvae. The Y-axis indicates the percentage of larval survival, and the X-axis shows the number of days following larval hatching. The experiment consisted of four trials, each with 25 larvae. **(E)** Graph representing the percentage of pupariation after egg laying (AEL) used to assess developmental time for control and *dOCRL^KO^*. Y-axis represents the percentage of larvae pupariated, and the X-axis shows the time in hours post egg laying. **(F)** Quantification of the average weight of third instar wandering larvae in control, *dOCRL^KO^* and *dOCRL^KO^;hs>HA::dOCRL*. The Y-axis indicates the weight, measured in milligrams, for five larvae per sample of each genotype (n=10) .Column plots with mean and standard deviation are shown. **(G)** The percentage of pupariation was quantified for control, *dOCRL^KO^* and *dOCRL^KO^;hs>HA::dOCRL*. The Y-axis indicates the proportion of larvae that underwent pupariation. Column plots with mean and standard deviation are shown. Each individual point on the graph represents a trial, with each trial comprising 25 larvae. Statistical tests: (A, F and G) Student’s unpaired t-test is used.****p<0.0001. (D) Survival curve used Long-rank (Mantel-cox) test and Gehan-Breslow-Wilcoxon test to analyze the statistical significance between two genotypes, ****p<0.0001.

**Supplementary figure 2: Nephrocyte specific *dOCRL* knockout phenocopies the defects of whole body *dOCRL^KO^* (A)** The percentage of pupariation was quantified for control and *dOCRL^N-KO^*. The Y-axis indicates the proportion of larvae that underwent pupariation. Column plots with mean and standard deviation are shown. Each individual point on the graph represents a trial, with each trial comprising 25 larvae. **(B)** The proportion of eclosed flies was measured for both control and *dOCRL^N-KO^*. The Y-axis represents the percentage of flies that successfully eclosed. Column plots with mean and standard deviation are shown. **(C)** Quantification of the average weight of third instar wandering larvae in control, *dOCRL^KO^* and *dOCRL^N-KO^*. The Y-axis indicates the weight, measured in milligrams, for five larvae per sample of each genotype (n=12). Column plots with mean and standard deviation are shown. **(D)** Confocal micrographs of 10kDa TMR-dextran uptake assay in control and nephrocyte specific *dOCRL^KO^* (using SNS GAL4). Scale bar: 10 μm. **(E)** Quantification of dextran taken up by the cell in control and *Sns>dOCRL^KO^*; Y-axis shows the Mean fluorescent intensity (MFI) of 10KDa TMR-Dextran in nephrocytes per unit area calculated using ImageJ. (N=9, n=90). Violin plots with mean and standard deviation are represented. **(F)** Confocal micrographs of 10KDa TMR-dextran uptake assay in control and salivary gland specific *dOCRL^KO^* (using AB1 GAL4). Scale bar: 10 μm**. (G)** Quantification of dextran taken up by the cell in control and *AB1>dOCRL^KO^*; Y-axis shows the Mean fluorescent intensity (MFI) of 10kDa TMR-Dextran in nephrocytes per unit area calculated using ImageJ (N=9, n=90). Violin plots with mean and standard deviation are represented. **(H)** Confocal micrographs of 10KDa TMR-dextran uptake assay in control and hemocyte/lymph gland specific *dOCRL^KO^* (using Hml GAL4). Scale bar: 10 μm. **(I)** Quantification of dextran taken up by the cell in control and *Hml>dOCRL^KO^*; Y-axis shows the Mean fluorescent intensity (MFI) of 10KDa TMR-Dextran in nephrocytes per unit area calculated using ImageJ. (N=9, n=90). Violin plots with mean and standard deviation are represented. Statistical tests: (A, B, C, E, G and I) Student’s unpaired t-test is used. ns-Non significance, ****p<0.0001, *-p<0.05.

**Supplementary figure 3: dOCRL regulates late-endolysosomal compartment (A)** Confocal micrographs showing immunostaining of control, *dOCRL^KO^*and *dOCRL^KO^; Dot>Rab7^CA^* nephrocytes using the lysosomal marker Cathepsin L. Scale bar: 10 μm**. (B)** Quantification of Cathepsin L staining in control, *dOCRL^KO^* and *dOCRL^KO^; Dot>Rab7^CA^*; Y-axis shows the Mean fluorescent intensity (MFI) of Cathepsin L in nephrocytes per unit area calculated using ImageJ (N=9, n=90). Violin plots with mean and standard deviation are represented. **(C)** Multiple sequence alignment analysis of dOCRL with human OCRL and INPP5B by Clustal Omega. The conserved amino acid residue 468-D (aspartic acid) in dOCRL is mutated to Glycine by site directed mutagenesis. **(D)** Sanger sequencing chromatogram showing the GAT to GGT nucleotide change within the phosphatase domain of the dOCRL in LTS::dOCRL::GFP transgene. **(E)** Immunoblots from larval fillets containing only nephrocytes and body wall are probed with antibody against dOCRL to confirm LTS::dOCRL::GFP and LTS::dOCRL::GFP(PD) expression in Control, *dOCRL^KO^*, *dOCRL^KO^;Dot>LTS::dOCRL::GFP* and *dOCRL^KO^;Dot>LTS::dOCRL::GFP(PD)*. Tubulin is used as a loading control. (M_R_: molecular weight in kDa). Statistical tests: (B) Student’s unpaired t-test is used, ****p<0.0001.

**Supplementary figure 4: Lysosomal-targeted dOCRL rescues early endosomal defects in *dOCRL^KO^* (A)** Confocal micrographs showing immunostaining of control, *dOCRL^KO^* and *dOCRL^KO^; Dot>LTS::dOCRL* nephrocytes using the early endosome marker Rabenosyn-5. Scale bar: 10 μm. **(B)** Quantification of Rabenosyn-5 staining in control, *dOCRL^KO^* and *dOCRL^KO^; Dot>LTS::dOCRL*; Y-axis represents the Mean fluorescent intensity (MFI) of Rabenosyn-5 in nephrocytes per unit area calculated using ImageJ (N=4, n=40). Violin plots with mean and standard deviation are represented. **(C)** Confocal micrographs of mBSA-Cy5 uptake assay in control, *dOCRL^KO^* and *dOCRL^KO^; Dot>LTS::dOCRL.* Scale bar: 10 μm. **(D)** Quantification of mBSA taken up by the cell in control, *dOCRL^KO^* and *dOCRL^KO^; Dot>LTS::dOCRL*; Y-axis shows the Mean fluorescent intensity (MFI) of mBSA-Cy5 in nephrocytes per unit area calculated using ImageJ (N=9, n=90). Violin plots with mean and standard deviation are represented. Statistical tests: (D) Student’s unpaired t-test is used. ****p<0.0001.

**Supplementary figure 5: Verification of uniform probe expression in nephrocytes of various genotypes used in this study (A)** Immunoblots from larval fillets containing only nephrocytes and body wall are probed with antibody against mCherry to confirm PH-PLCδ::mCherry expression in Control and *dOCRL^KO^*. Syntaxin-1a is used as a loading control. (M_R_: molecular weight in kDa). **(B)** Quantification of PH-PLCδ::mCherry probe levels in Control and *dOCRL^KO^*. Column plots with mean and standard deviation are represented. **(C)** Immunoblots from larval fillets containing only nephrocytes and body wall are probed with antibody against GFP to confirm P4M::GFP expression in Control and *dOCRL^KO^*. Syntaxin-1a is used as a loading control. (M_R_: molecular weight in kDa). **(D)** Quantification of P4M::GFP probe levels in Control and *dOCRL^KO^*. Column plots with mean and standard deviation are represented. **(E)** Immunoblots from larval fillets containing only nephrocytes and body wall are probed with antibody against mCherry to confirm mCherry::2XFYVE expression in Control and *dOCRL^KO^*. Syntaxin-1a is used as a loading control. (M_R_: molecular weight in kDa). **(F)** Quantification of mCherry::2XFYVE probe levels in Control and *dOCRL^KO^*. Column plots with mean and standard deviation are represented. **(G)** Immunoblots from larval fillet containing only nephrocytes are probed with antibody against mCherry to confirm LTS::mCherry expression in Control and *dOCRL^KO^*. Tubulin is used as a loading control. (M_R_: molecular weight in kDa). **(H)** Quantification of LTS::mCherry probe levels in Control and *dOCRL^KO^*. Column plots with mean and standard deviation are represented. Statistical tests: (B, D, F and H) Student’s unpaired t-test is used. ns-non significance.

**Supplementary Table 1:** The table presents the candidate genes evaluated in the genetic screen. A “+” in the uptake column signifies that AgNO_3_ was successfully taken up by nephrocytes, whereas a “–” indicates a complete absence of AgNO_3_ uptake in nephrocytes. In the clearance column, a “+” denotes efficient clearance of AgNO_3_ by nephrocytes, while a “–” denotes reduced clearance. The designation “N.E.” indicates that these cells were not evaluated due to compromised uptake.

