## Supplementary Figures 1-5 for "OCRL regulates lysosomal function and endolysosomal homeostasis in *Drosophila* nephrocytes"

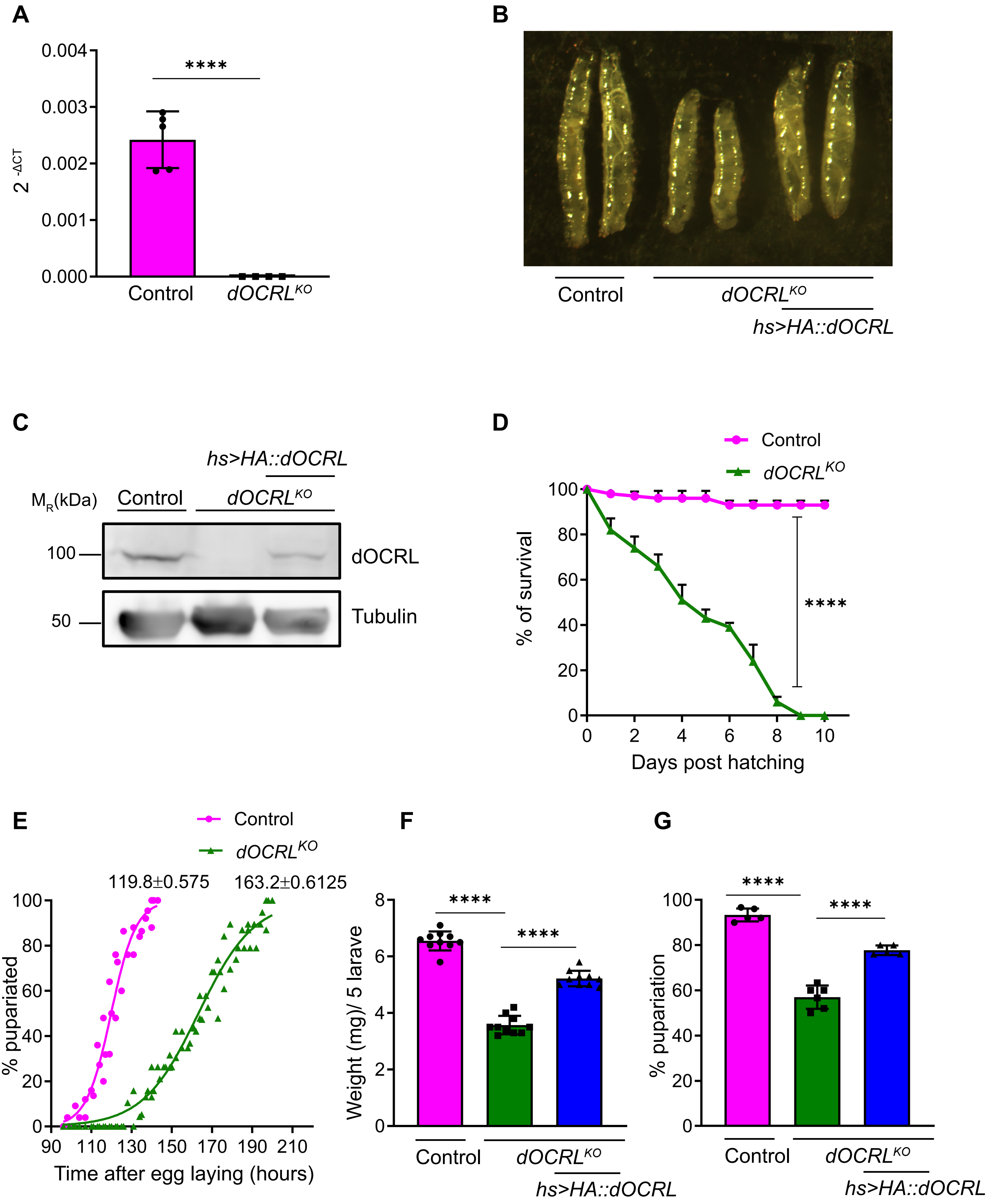


**Supplementary figure 1**

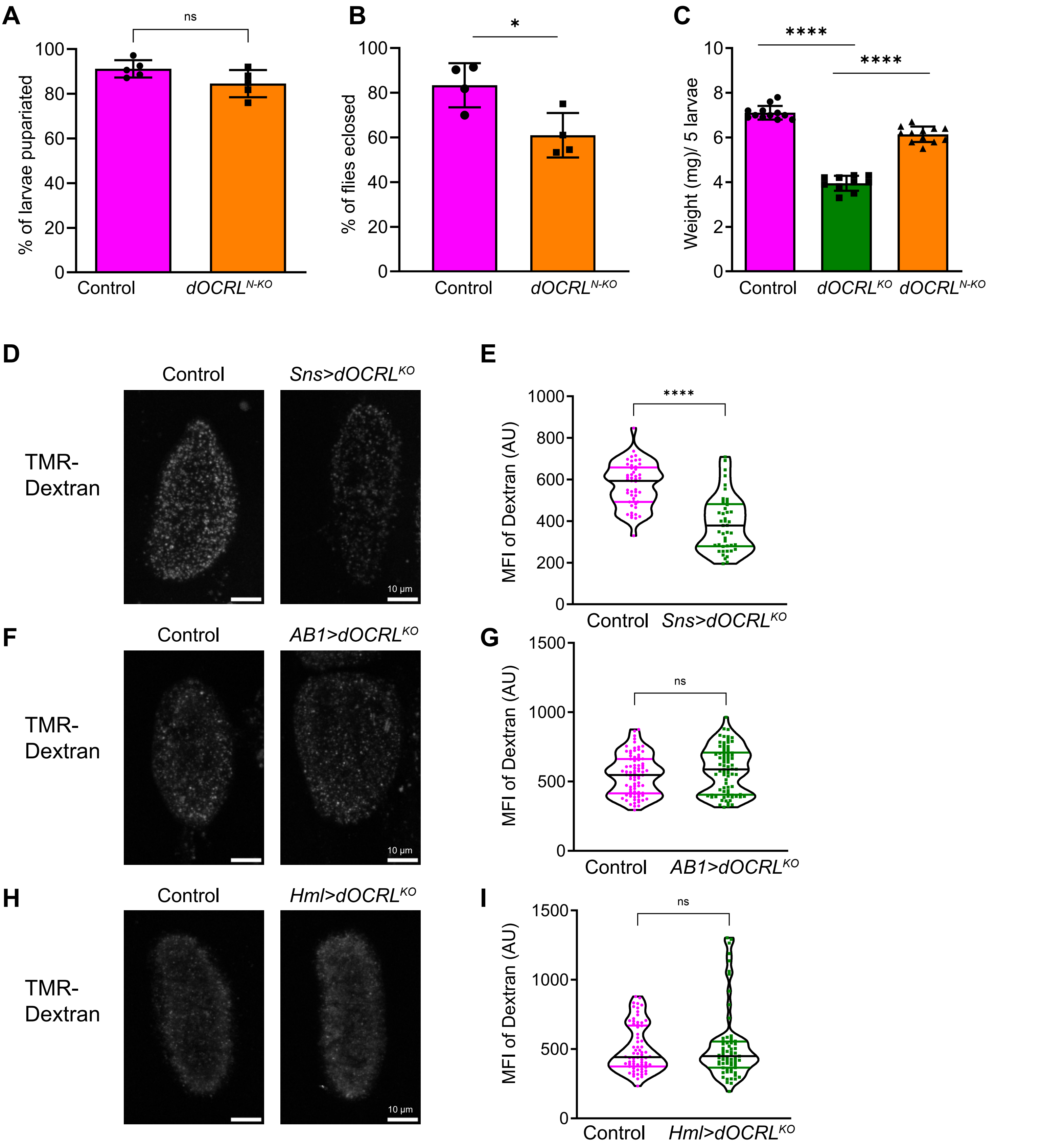


**Supplementary figure 2**

**A**

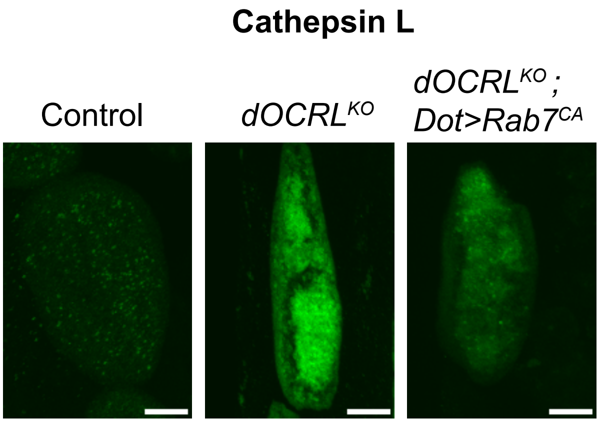


**B**

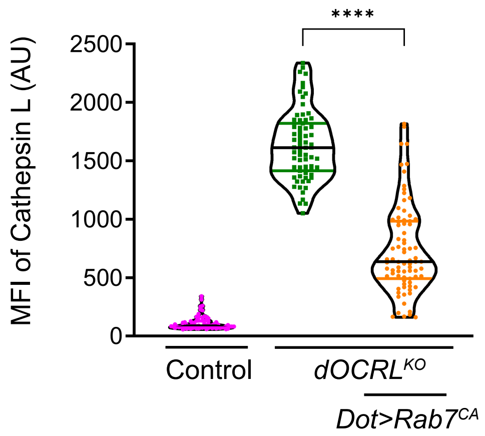


**C**

```
dOCRL
KFRPTYKYDPGTDNYDSSEKQRAPAYCDRVLWKGTRIEQLAYNSIMEIRQSDHKPVYAVF     476
hOCRL
KFIPTYKYDSKTDRWDSSGKCRVPAWCDRILWRGTNVNQLNYRSHMELKTSDHKPVSALF     531
hINPP5B
TFQPTYKYDTGSDDWDTSEKCRAPAWCDRILWKGKNITQLSYQSHMALKTSDHKPVSSVF     556
                  .* ******  :* :*:* * *.**:***:**:*..: ** *.* * ::
****** ::*
```

**D**

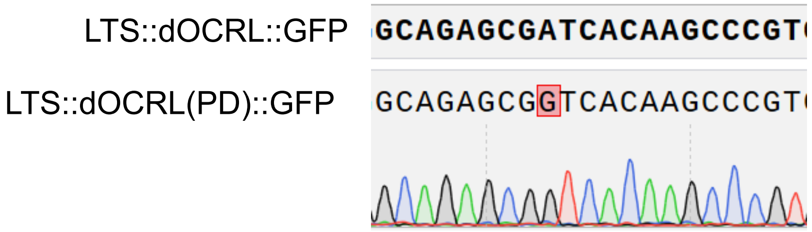


**E**

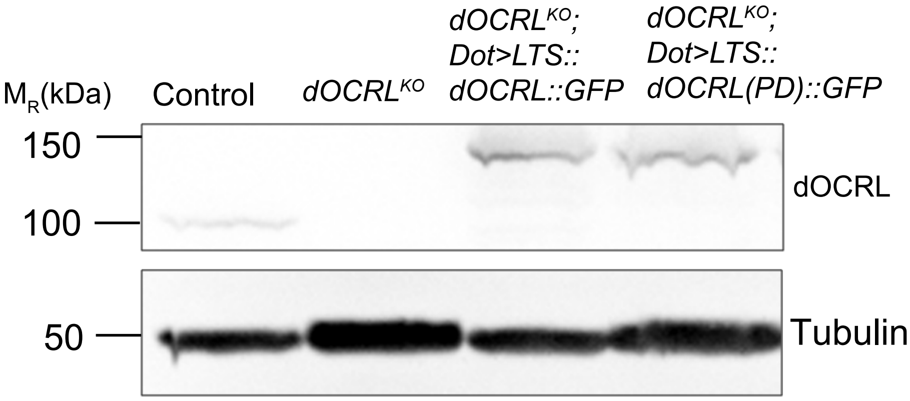


**Supplementary figure 3**

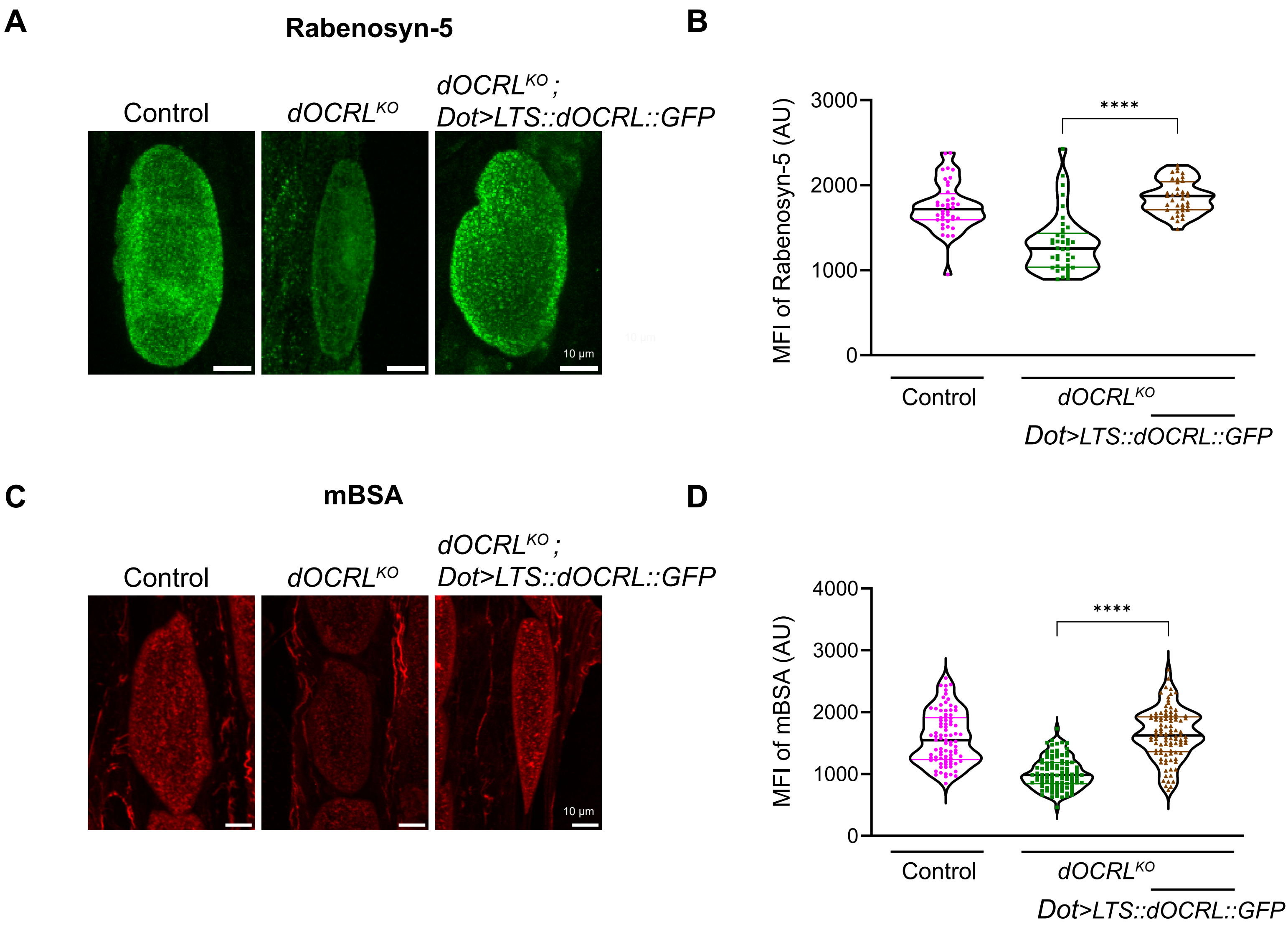


**Supplementary figure 4**

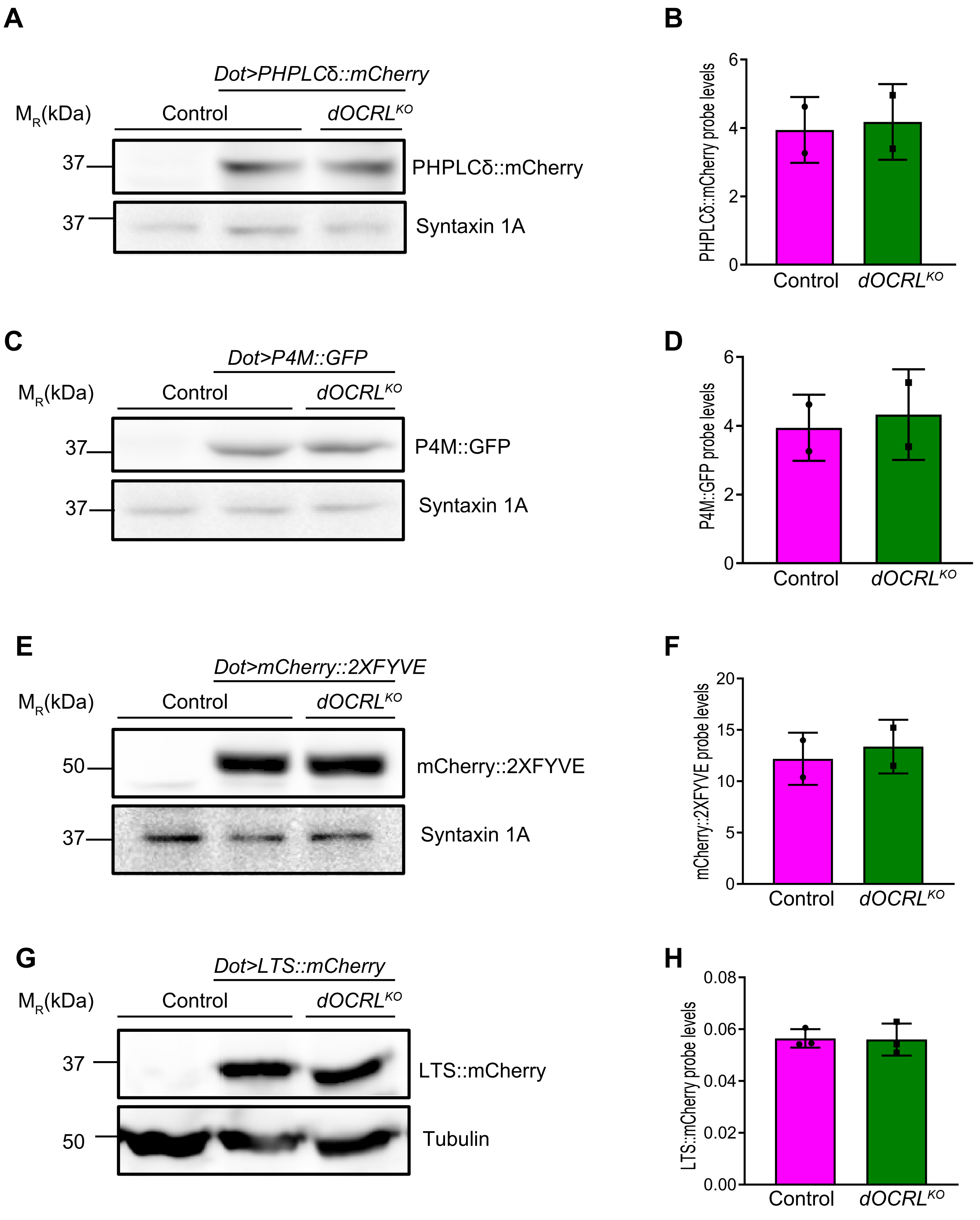


**Supplementary figure 5**
