## Supplementary Table 1 for "OCRL regulates lysosomal function and endolysosomal homeostasis in *Drosophila* nephrocytes"

| Candidate genes | Nature of allele | Uptake | Clearance |
|---|---|---|---|
| Wild-type | | + | + |
| *dOCRL* | Knock-out | + | - |
| *Rab7* | Dominant-negative | + | - |
| *Shibire* | RNAi | - | N.E. |
| *Rab5* | Dominant-negative | - | N.E. |
| *ClC-c (Clcn-5)* | RNAi | - | N.E. |
| *Rab11* | Dominant-negative | - | N.E. |
| *ATG1* | Over-expression | - | N.E. |
| *Cathepsin L* | Knock-out | - | N.E. |
| *Trpml* | Knock-out (null allele) | - | N.E. |
| *Trpml* | RNAi | + | + |
| *Rab4* | Dominant-negative | + | + |
| *Rab27* | Dominant-negative | + | + |
| *Rab2* | Dominant-negative | + | + |
| *Syntaxin 17* | RNAi | + | + |
| *ClC-b (Clcn-7)* | Knock-out (null allele) | + | + |
| *Vha39* | RNAi | + | + |
| *Fab1* | RNAi | + | + |

**Supplementary Table 1:** The table presents the candidate genes evaluated in the genetic screen. A "+" in the uptake column signifies that $AgNO_3$ was successfully taken up by nephrocytes, whereas a "–" indicates a complete absence of $AgNO_3$ uptake in nephrocytes. In the clearance column, a "+" denotes efficient clearance of $AgNO_3$ by nephrocytes, while a "–" denotes reduced clearance. The designation "N.E." indicates that these cells were not evaluated due to compromised uptake.
